# AbSolution: interactive exploration of sequence-derived features in AIRR-seq repertoires

**DOI:** 10.64898/2026.05.20.726477

**Authors:** Rodrigo García-Valiente, Charisios Triantafyllou, Barbera van Schaik, Aldo Jongejan, Sabrina Pollastro, Dornatien Anang, Jeroen E. J. Guikema, Niek de Vries, Huub Hoefsloot, Antoine H.C. van Kampen

## Abstract

High-throughput sequencing of B-cell and T-cell immune receptor repertoires provides unprecedented insight into adaptive immune responses. The data produced are structured by clonal relationships and somatic mutation signatures, and yield extremely rich information in sequence-derived features, including physicochemical properties and compositional patterns. However, integrated analysis across datasets, conditions, and time points remains challenging. Current analytical tools typically focus only on certain features within individual repertoires, without enabling integrated, multivariable comparisons across datasets, conditions, and time points to address their diversity and variability. Here we present AbSolution, a user-friendly and flexible interactive application for comprehensive exploration of immune repertoires and their sequence-based properties. AbSolution enables multiscale analysis of thousands of sequence-derived features across receptor regions, while accounting for V(D)J usage, clonal composition and experimental groupings. We demonstrate its utility by identifying distinct sequence-based profiles associated with dominant (highly abundant) and non-dominant B-cell clones in peripheral blood BCR repertoires from patients with idiopathic inflammatory myopathies, and with antigen-responsive T-cell populations over time in a longitudinal *in vitro* antigen-stimulation dataset. Through interactive, interlinked visualizations, statistical feature selection and multi-sample comparisons, AbSolution facilitates integrated feature profiling that supports the interpretation of immune selection processes and enables systematic analysis of complex repertoire datasets.

## Introduction

T-cell receptors (TCRs) are heterodimeric proteins, canonically composed of either an alpha and a beta chain (αβ TCRs) or a gamma and a delta chain (γδ TCRs) (Brenner et al., 1986; Hochstenbach & Brenner, 1989; Kreslavsky et al., 2010; Saito et al., 1984). B-cell receptors (BCRs) contain a membrane-bound immunoglobulin (Ig) heterotetrameric protein, composed of two heavy chains (encoded by the IGH locus) and two κ or λ light chains (encoded by the IGK and IGL loci, respectively); antibodies (Abs) are the soluble form of the Ig (Alt et al., 1980; Dong et al., 2022; Edelman & Poulik, 1961; Porter, 1959). Each TCR and BCR chain harbours three complementarity-determining regions (CDR1-3), representing the most variable parts of the receptor which determine a large part of the interaction with the antigen (Ag). Four framework regions (FWR1-4) primarily provide structural support to the receptor (Lefranc et al., 2003; Sela-Culang et al., 2013; Wu & Kabat, 1970). Both CDRs and FWRs are located within the variable domain of the receptor chains (Edelman et al., 1969; Lefranc et al., 2003).

The generative potential of unique BCRs and TCRs in humans and other mammals is immense. In humans, the naïve B-cell repertoire diversity has been estimated to include up to 10^18^ different variants (Rees, 2020). The potential human T-cell repertoire diversity before thymic selection has been estimated to include up to 10^15^ variants (Nikolich-Zugich et al., 2004). This TCR and BCR diversity arises through V(D)J somatic recombination in individual lymphocytes, a stochastic but biased process in which V, D and J gene segments are joined in Ig heavy and TCR β/δ chains, and V and J segments in Ig light and TCR α/γ chains (Hozumi & Tonegawa, 1976; Yancopoulos et al., 1984). Additional diversity is generated at the recombination junctions through imprecise joining, exonuclease-mediated nucleotide trimming, palindromic (P) nucleotide additions and terminal deoxynucleotidyl transferase (TdT)-mediated untemplated nucleotide additions, collectively referred to as junctional diversity (Alt & Baltimore, 1982; Desiderio et al., 1984; Lafaille et al., 1989). This diversity is increased through the pairing of the corresponding chains (Dupic et al., 2019; Schroeder & Cavacini, 2010). In antigen-experienced B cells, BCR sequences can be further diversified by the process of somatic hypermutation (SHM) in the germinal centre (GC) (Peled et al., 2008).

The full repertoire of B- and T-cell receptors can be explored through adaptive immune receptor repertoire sequencing (AIRR-seq) (Boyd & Crowe, 2016; Klarenbeek et al., 2010; Liu et al., 2021). Using AIRR-seq data, receptor sequences can be reconstructed and compared to their inferred germline sequences. Sequences derived from the same original rearrangement event are considered part of the same clone, reflecting descent from a common ancestor. Multiple computational strategies have been proposed to cluster receptor sequences into putative clones based on shared sequence features, including hierarchical distance-based methods (Gupta et al., 2017). Such clones represent lineages, and their relative frequency is determined by the number of receptor sequences assigned to each clone divided by the total number of sequenced receptors. Analysis of this data holds much clinical and translational potential (Arnaout et al., 2021) such as the monitoring of immune responses during (auto)immune disease (Anang et al., 2023; Bashford-Rogers et al., 2019; Pollastro et al., 2024), and the determination of tumour infiltrating lymphocytes (Raphael et al., 2025). AIRR-seq has also been used for the development of monoclonal Ab therapeutics (Goldstein et al., 2019; Setliff et al., 2019) and vaccines (Roskin et al., 2020; Taft et al., 2022).

Clonal frequencies and V(D)J gene usage capture only part of the information contained in immune receptor repertoires. Sequence-derived properties, including physicochemical characteristics, compositional patterns and mutational features, can provide additional insight into receptor structure, clonal selection and antigen recognition. A growing body of evidence has demonstrated that such features are associated with functional and disease-associated repertoire characteristics. For instance, amino acid physicochemical properties of CDR3 regions distinguish different subsets within T cells (Kasatskaya et al., 2020) and B cells (Ghraichy et al., 2021). Amino acid descriptor sets (Todeschini & Consonni, 2000), including reduced-dimensional representations such as Kidera factors (Kidera et al., 1985), FASGAI vectors (Liang et al., 2008) or BLOSUM indices (Georgiev, 2009), have been shown to capture functionally relevant BCR variation linked to antigen recognition (Laffy et al., 2017), to disease type (Crescioli et al., 2023) and to Ig chain type (Townsend et al., 2016). Identical representations have been used to model (Shcherbinin et al., 2023) and predict (Tatikonda et al., 2024) TCR-epitope specificity. Finally, V-gene-specific amino acid substitution biases resulting from SHM exhibit partial correlation with underlying physicochemical properties (Sheng et al., 2017), indicating context-dependent structural changes.

Despite advances in feature-based repertoire analysis, systematic and integrative investigation of sequence-derived properties remains challenging. Numerous features can be computed using existing packages and computational resources, including the R packages *alakazam* (Gupta et al., 2015) and *Peptides* (Osorio et al., 2015), the repertoire analysis framework VDJtools (Shugay et al., 2015), and databases such as AAindex (Kawashima & Kanehisa, 2000). However, these approaches typically require programming expertise and additional downstream analyses to identify significant features. In addition, their scope varies widely, often being restricted to specific sets of physicochemical properties or receptor regions. In parallel, several dedicated applications have been developed for sequence-based feature exploration and analysis. In the case of B-cell repertoires, BRepertoire (Margreitter et al., 2018) enables the study of different CDR3 physicochemical properties and clonotype clustering, whereas AIRRscape (Waltari et al., 2022) allows SHM-level analysis and clonal topology exploration. In the case of T-cell repertoires, GENTLE (Andrade et al., 2023) supports the identification of differentially represented k-mers (2- to 4-amino-acid length). While all of these applications enable exploration of specific properties in some regions and repertoires, they are limited in their ability to integrate different levels of repertoire information, perform downstream sequence-level analysis, or facilitate direct comparison of features across regions, samples, time points, or experimental conditions.

To address these limitations, we developed AbSolution, an interactive R Shiny application for comprehensive exploration and analysis of sequence-derived features in adaptive immune receptor repertoires. AbSolution integrates large-scale computation of physicochemical properties, amino acid descriptor sets, sequence motifs, compositional patterns, and SHM-related features across receptor regions and complete variable domains. By analysing both the observed repertoire sequences and their reconstructed germline counterparts, AbSolution enables quantitative assessment of selection-associated sequence evolution. The application supports multiscale stratification of repertoires according to samples, experimental groups, clonal classification, dominance patterns, VJ usage, isotypes, and temporal dynamics. Through interactive visualizations, feature-wise statistical testing, configurable normalization and subsampling strategies, and automated export of code, figures, and metadata, AbSolution enables reproducible, hypothesis-driven and discovery-oriented analyses while controlling for common confounding factors. Here, we present the design and implementation of AbSolution and demonstrate its utility using TCR and BCR repertoire datasets, revealing sequence-derived profiles associated with antigen-driven expansion, clonal persistence and clonal dominance.

## Results

To illustrate the application of AbSolution, we present two use cases: one based on a longitudinal TCR repertoire dataset during antigen stimulation and another based on a BCR repertoire dataset from patients with idiopathic inflammatory myopathies.

### Antigen-responsive T-cell clones in longitudinal samples display distinct physicochemical TCR properties

We used AbSolution to perform a feature-based comparison of two samples from a TCR β-chain AIRR-seq dataset adquired from peripheral blood mononuclear cells of a single healthy donor before (day 0) and after (day 10) *in vitro* antigen stimulation culture with a mix of known CD4+ T-cell epitopes derived from cytomegalovirus, Epstein-Barr virus, and influenza A virus (Pollastro et al., 2020).

Within the application, as the TCR dataset was processed without UMI-consensus-based PCR-error correction, we applied a conservative downstream quality filter and excluded sequences with more than three substitutions (putative sequencing-error events) predicted to alter the amino acid sequence. The analysis presented here was restricted to sequence length and (composite) features describing amino-acid physicochemical properties, computed across the complete variable sequence and individual receptor regions. Of these 1368 features, a total of 734 showed non-zero variance across the sequences and were included in the downstream analysis. Their distribution across regions is summarized in *Table 1*. Statistically significant features between groups were identified using feature-wise logistic regression with Benjamini-Yekutieli multiple testing correction (Benjamini & Yekutieli, 2005).

**Table 1.** Number of sequence-derived features per region identified across different TCR repertoire comparisons. Values indicate the total number of computed features and the subset of features identified as statistically significant following feature-wise analysis for (i) population-level comparison between day 0 and day 10, (ii) comparison between shared and unique clones at day 0 (early and persistent clones) versus unique clones at day 10 (late-detected clones), and (iii) comparison between differentially under-expanded (DUE) and differentially over-expanded (DOE) clones. Statistical significance was determined using feature-wise logistic regression with multiple testing correction (Benjamini-Yekutieli). FWR4 contains fewer computed features due to the exclusion of N-glycosylation motif-related features, which showed zero variance in this region.

| Region | Computed features | Statistically significant features (day 0 vs day 10) | Statistically significant features (early and persistent clones vs late-detected clones) | Statistically significant features (DUE vs DOE clones) |
| --- | --- | --- | --- | --- |
| **FWR1** | 92 | 59 | 57 | 71 |
| **CDR1** | 92 | 43 | 49 | 75 |
| **FWR2** | 92 | 53 | 59 | 71 |
| **CDR2** | 92 | 43 | 30 | 67 |
| **FWR3** | 92 | 63 | 63 | 75 |
| **CDR3** | 92 | 43 | 46 | 70 |
| **FWR4** | 90 | 57 | 51 | 74 |
| **Variable** | 92 | 61 | 61 | 74 |

Subsequently, we analysed these longitudinal TCR repertoire dynamics using three complementary stratifications: population-level comparisons between day 0 and day 10 (Figures 1 A-C), contrasts between pre-existing or persistent clones and newly detected clones (Figures 1 D-F), and classification of clones according to their expansion behaviour following antigen stimulation (Figures 1 G-I). Within AbSolution, these figures are dynamic and linked: selecting sequences, clones or clonal intersections in one panel updates the corresponding information across other visualizations and input parameters, enabling direct exploration of feature distributions and retrieval of the underlying sequences for downstream analyses.

**Figure 1.**
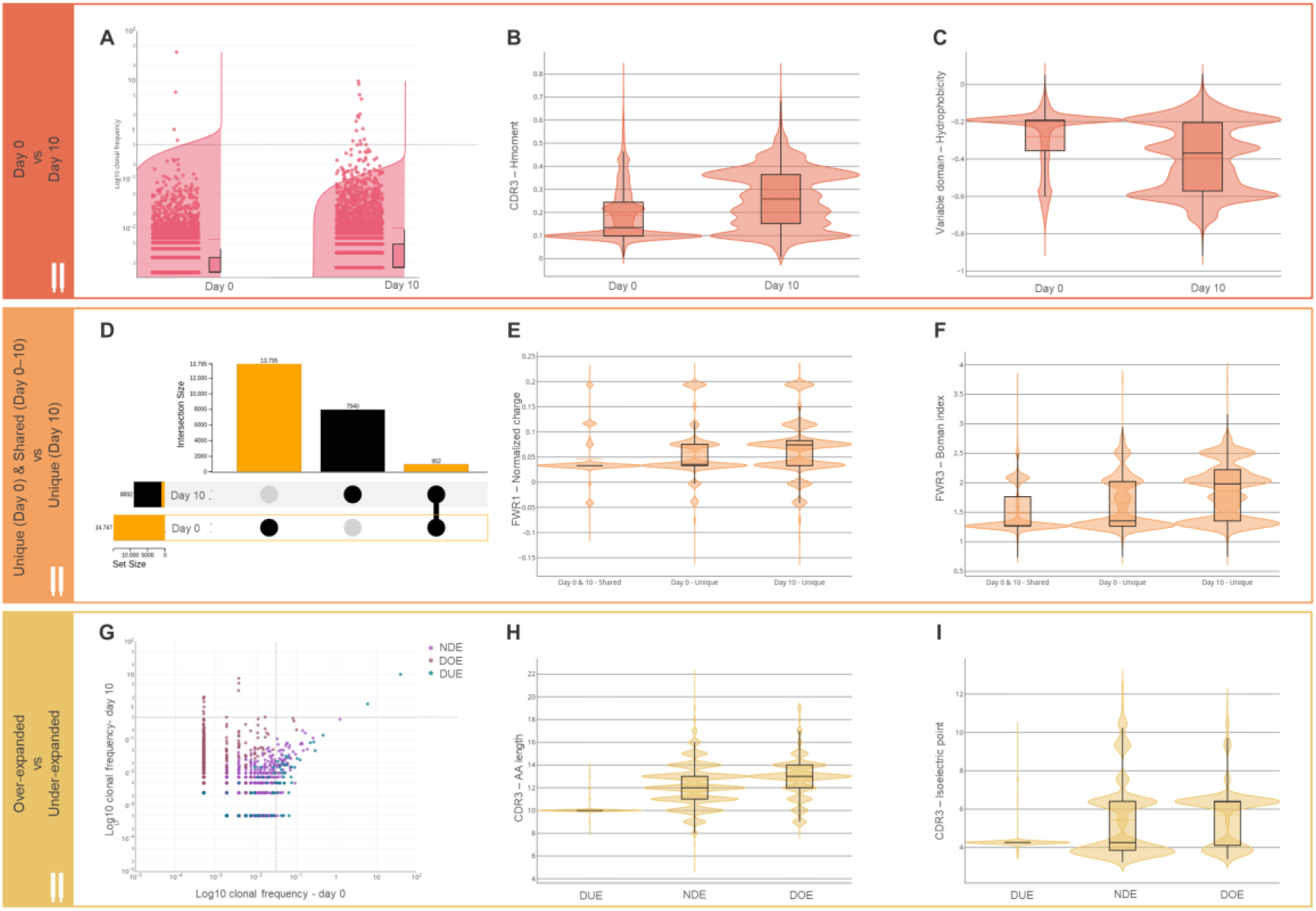
Longitudinal analysis of T-cell clones and selected statistically significant physicochemical features at day 0 and day 10 after antigen stimulation. (A-C) Comparison of clone populations at days 0 and 10. **(A)** Split-violin plots with box-plots showing the relative frequency distribution (log₁₀-transformed) of TCR clone sizes at day 0 and day 10 following antigen stimulation. Each dot represents an individual clone, and clonal groups were defined within AbSolution based on V(D)J usage and identical CDR3 nucleotide sequence. The violin plots are split to allow visualization of clones stratified according to their germline sequence, as illustrated in Figure 2B-E,G. The dashed horizontal line represents the threshold for clonal dominance, 0.5% (Klarenbeek et al., 2012). **(B)** Violin plots of CDR3 hydrophobic moment at day 0 and day 10. **(C)** Violin plots of variable domain hydrophobicity at day 0 and day 10. **(D-F) Comparison of shared and unique clones at day 0 versus unique clones at day 10. (D)** UpSet plot summarizing clone intersections between day 0 and day 10, highlighting (orange) clones unique to day 0 and shared between day 0 and day 10. **(E)** Violin plots of normalized-by-length FWR1 charge for clones shared between time points, unique to day 0, and unique to day 10. **(F)** Violin plots of FWR3 Boman index for clones shared between time points, unique to day 0, and unique to day 10. **(G-I) Comparison of differentially over-expanded (DOE) and under-expanded clones (DUE) at day 10.** Clones are coloured as DOE, DUE or not-differentially expanded (NDE) clones according to the original study, based on their relative abundance changes from day 0 to day 10 post-stimulation. NDE were excluded from feature discrimination analyses but are shown in the visualizations for completeness. **(G**) Scatter plot comparing log₁₀-transformed clonal frequencies at day 0 and day 10. Dashed lines represent the clonal dominance threshold (0.5%). **(H)** Violin plots of CDR3 amino acid length across expansion categories (DUE, NDE, DOE). **(I)** Violin plots of CDR3 isoelectric point values across expansion categories. Boxplots embedded within violin plots indicate medians and interquartile ranges.

To compare population-level differences between day 0 and day 10, sequences from both time points were grouped into clones within AbSolution based on shared V(D)J gene usage and identical CDR3 nucleotide sequences. *Figure 1A* shows a marked post-stimulation increase in the number of dominant clones (frequency ≥ 0.5% (Klarenbeek et al., 2012)), from 4 on day 0 to 14 on day 10. A total of 422 features were identified as statistically significant, spanning both the complete variable domain and its individual regions (*Table 1*). Among others, feature analysis with AbSolution further showed that this expansion is accompanied by a statistically significant increase in the hydrophobic moment, a measure of the spatial segregation of hydrophobic and hydrophilic residues (Eisenberg et al., 1982), of CDR3 (*Figure 1B*) and a statistically significant decrease in overall hydrophobicity of the variable domain (Kyte & Doolittle, 1982) (*Figure 1C*). These observations suggest that expanded clones display CDR3 regions with a more pronounced amphipathic organization embedded within a more polar variable domain that facilitates structured interaction surfaces (Tsuchiya et al., 2018).

Next, AbSolution was used to compare early and persistent clones with late-detected clones (*Figure 1D*). A total of 416 features were identified as statistically significant, distributed across the full variable domain and individual regions (*Table 1*). In this dataset, clones unique to day 10 compared to day 0 and to the clones present at both time points showed both increased FWR1 normalized charge (*Figure 1E*) and FWR3 Boman index (*Figure 1F*), which predicts the protein-binding potential of a peptide (Boman, 2003). The increased FWR3 Boman index may be related to the HV4 loop, that through binding can contribute to ligand recognition and modulate interactions with peptide-MHC complexes (Michielin et al., 2000). Because FWRs provide the structural scaffold that determines the positioning and orientation of antigen-binding loops, differences in their physicochemical properties may influence how the TCR presents its peptide-MHC interaction surface. These observations suggest that emerging antigen-responsive T cells are associated with certain FWR characteristics to support structural positioning of peptide-MHC interfaces (Rosenberg et al., 2024).

Finally, we used AbSolution to characterize sequences according to their expansion behaviour following antigen stimulation. For this analysis, we considered differentially over-expanded (DOE), differentially under-expanded (DUE), and not differentially expanded (NDE) clones, as defined in the original publication (Pollastro et al., 2020). Clones in these subsets were defined based on a statistically significant change in their frequency at day 10 compared to day 0, and a log_2_ fold change ≥1.5 (DOE), ≤-1.5 (DUE), or between -1.5 and 1.5 (NDE). *Figure 1G* shows these three subgroups in the context of clonal frequencies. A total of 577 features were identified as statistically significant, with broad representation across both the full variable domain and individual regions (*Table 1*). Compared to DUE clones, DOE clones showed both statistically significant increased CDR3 length (*Figure 1H*) and CDR3 isoelectric point (*Figure 1I*). We did not compare DOE/DUE to NDE clones to determine significance for these features, but did include the values for NDE clones in the figure to show that these represent an intermediate state between DOE and DUE. The CDR3 region mediates most contacts with the peptide–MHC complex, and these features may enhance structural reach and electrostatic complementarity. Therefore, the enrichment of longer and more positively charged CDR3 sequences suggests selective expansion of clones with favourable antigen-recognition properties during stimulation, which is consistent with affinity-driven clonal expansion of specific TCRs (Busch & Pamer, 1999).

Taken together, these analyses demonstrate how AbSolution is used to explore and reveal antigen-driven changes in T-cell clonal abundance associated with trends in TCR physicochemical properties. AbSolution uncovers complementary sets of informative features across multiple levels of repertoire organization, highlighting the value of flexible stratification strategies for interpreting immune repertoire dynamics.

### Identification of dominant B-cell clones and characterization of their distinct sequence-based properties

As a second use case, we applied AbSolution to analyse six samples from a BCR IgH AIRR-seq dataset from peripheral blood of patients with idiopathic inflammatory myopathies (myositis) (Anang et al., 2023). Clonal abundance, and therefore, dominance, is a result of Ag-driven clonal expansion and selection in GCs. The objective of the analysis is to identify sequence features that distinguish dominant (positively selected, expanded) B-cell clones from non-dominant clones, and to determine whether these differences are already present at the germline level or arise through SHM and clonal selection.

Within AbSolution, we first selected sequences containing between 0 and 25 replacement mutations relative to their germline counterparts to retain unmutated to moderately mutated sequences and exclude highly mutated outliers (Weill & Reynaud, 2020). Following earlier definitions, within the application we defined clones with a frequency greater than or equal to 0.5% of the analysed BCR sequences to be dominant clones (Klarenbeek et al., 2012). We focused exclusively on features related to sequence length, N-glycosylation sites, hot-spot and cold-spot motifs, peptide properties, and replacement/silent (R/S) mutation-related features for individual regions and the full sequence. Of these 928 features, a total of 854 showed non-zero variance and were therefore included. Their distribution across regions and analysis stratifications (all samples, one individual sample, and one VJ combination within one individual sample) is summarized in *Table 2*. FWR1 was excluded from the analysis as the region was only partially sequenced. For each comparison, statistically significant features between clone types were identified using feature-wise logistic regression with Benjamini-Yekutieli multiple testing correction (Benjamini & Yekutieli, 2005).

**Table 2.** Number of sequence-derived features per region across different BCR analysis scopes. Values indicate the total number of computed features and the subset of features identified as statistically significant following feature-wise analysis at the level of all samples, a single sample (S15), and a restricted subset of sequences sharing the same VJ combination (IGHV3-15*01-IGHJ4*02). FWR1 was excluded from the analysis as the region was only partially sequenced. Statistical significance was determined using feature-wise logistic regression with multiple testing correction (Benjamini-Yekutieli).

| Region | Computed features | Statistically significant features (all samples) | Statistically significant features (sample S15) | Statistically significant features (S15, IGHV3-15*01-IGHJ4*02) |
| --- | --- | --- | --- | --- |
| CDR1 | 122 | 37 | 67 | 0 |
| FWR2 | 122 | 63 | 60 | 0 |
| CDR2 | 122 | 43 | 57 | 0 |
| FWR3 | 122 | 65 | 75 | 0 |
| CDR3 | 122 | 33 | 66 | 80 |
| FWR4 | 122 | 59 | 61 | 0 |
| Variable | 122 | 68 | 67 | 0 |

We compared the features of dominant and non-dominant clones across samples using logistic regression. Dominant clones were present in all samples (*Figure 2A*). When pooling data across patients, a total of 368 features were statistically significant. These features were identified across both the complete variable sequence and individual regions, with region-specific variation in feature number (*Table 2*). Dominant clones exhibited statistically significant increased FWR2 hydrophobicity (*Figure 2B*) and statistically significant decreased CDR3 normalized charge (*Figure 2C*) relative to non-dominant clones. Notably, FWR2 hydrophobicity in dominant clones differed significantly from the corresponding hydrophobicity of the reconstructed germline sequences (*Figure 2B*). Together, these results show that dominant clones differ from non-dominant clones in region-specific biophysical features, including, within this patient cohort, increased FWR2 hydrophobicity and reduced CDR3 normalized charge. Comparison with reconstructed germline sequences further indicates that these profiles do not arise from a single source: some differences reflect baseline sequence characteristics, whereas others are consistent with SHM-associated changes acquired during clonal expansion. This illustrates how AbSolution can distinguish repertoire-level feature differences from germline-associated effects and mutation-associated shifts.

**Figure 2.**
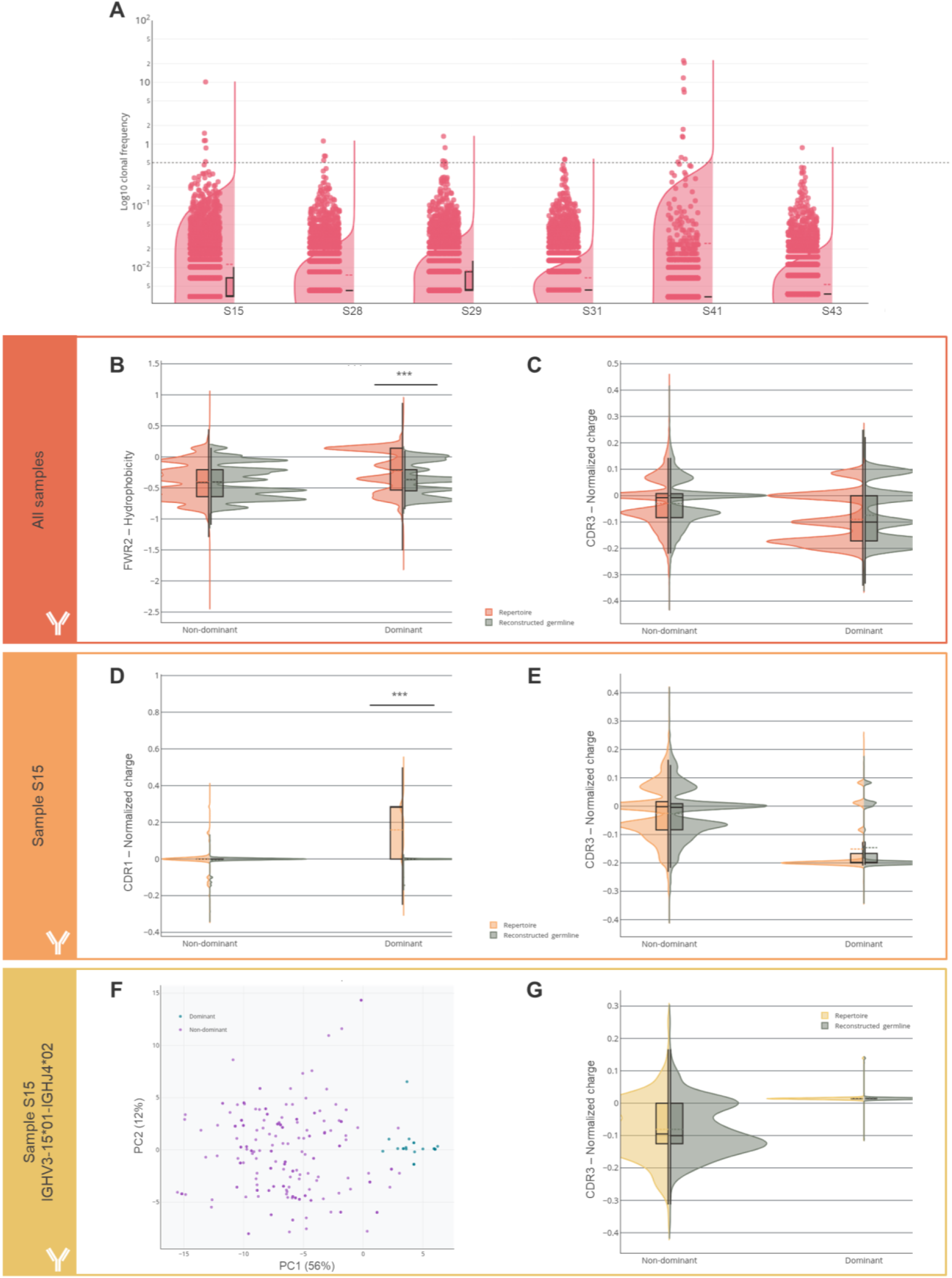
Identification of dominant B-cell clones and visualization of several significant physicochemical profiles. **(A)** Split violin and box plots showing log₁₀-transformed clonal frequencies across six myositis patient samples. Each dot represents an individual clone; the dashed horizontal line indicates the threshold used to classify dominant versus non-dominant clones. According to this threshold, the six samples present, respectively, 6, 4, 4, 2, 11 and 1 dominant clones. **(B, C)** Split-violin plots comparing FWR2 peptide hydrophobicity and **(B)** CDR3 normalized-per-length charge **(C)** between dominant and non-dominant clones across all six patients. **(D, E)** Same comparisons shown for a representative sample (S15) focusing on CDR1 normalized-per-length charge **(D)** and CDR3 normalized-per-length charge **(E)**. **(F)** Principal Component Analysis visualization of sequences from sample S15 sharing the IGHV3-1501–IGHJ402 gene combination, based on all statistically significant features, with points coloured according to clonal dominance status. The plot shows separation between dominant and non-dominant clones. **(G**) Violin plot of CDR3 normalized-per-length charge values for dominant and non-dominant clones restricted to a single VJ combination (IGHV3-15*01 IGHJ4*02) in sample S15. For all violin plots, repertoire-derived sequences are shown in colour and reconstructed germline sequences in grey. In addition to these features being statistically significant between dominant and non-dominant clones, statistical significance of paired t-tests comparing repertoire and germline values was calculated and is shown when the effect size is small to large (Cohen’s d ≥ 0.2). Significance levels are indicated as follows: P < 0.05 (*), P < 0.01 (**), and P ≤ 0.001 (***).

Features identified based on all samples may not be statistically significant when considering only an individual sample. Vice versa, in a single sample we may identify features that are not significant when all samples are analysed together. For example, focusing on sample S15, a total of 453 features were statistically significant (*Table 2*), distributed across the complete variable domain and individual Ig regions. Results show a strong significant increase in CDR1 normalized charge in dominant clones acquired through SHM (*Figure 2D*), which is not significant when all samples are analysed together. In addition, this comparison also shows a statistically significant decrease in CDR3 normalized charge, yet with a more pronounced effect (*Figure 2E*). By contrast, FWR2 hydrophobicity, which is statistically significant in the analysis of all samples, is also among the significant features in the S15-specific analysis.

Even within individual samples, substantial heterogeneity exists within groups. Directly comparing sequences with the same VJ combinations helps identify key differences that are otherwise obscured when considering all VJ combinations simultaneously (Marcou et al., 2018). As an example, we examined sequences with the IGHV3-1501/IGHJ402 gene usage in sample S15, as this VJ combination consists of a similar number of dominant (n = 332) and non-dominant (n = 338) sequences. A total of 80 features were significant, all belonging to the CDR3 region (*Table 2*). Within AbSolution, we performed and visualized the sequences using principal component analysis (PCA) to assess whether the statistically significant features support selection-driven segregation. This showed a clear separation between dominant and non-dominant clones (*Figure 2F*), illustrating how AbSolution can identify feature differences between dominant and non-dominant sequences after controlling for VJ gene usage, with the remaining variation within this matched VJ background mainly captured by CDR3-derived features. Moreover, distinct clonotypes within the dominant clone, defined by unique variable region nucleotide sequences (Sofou et al., 2023), occupy partially separated regions of the PCA feature space, representing functional heterogeneity within the clone. The dominant clone shows, in contrast to the group- and sample-level trends, elevated CDR3 normalized charge compared to their non-dominant counterparts (*Figure GF*).

Together, these results demonstrate that dominant B-cell clones exhibit reproducible physicochemical signatures across multiple analytical levels, including groups, samples, sequences with shared V(D)J gene usage, and clones. By resolving these patterns across multiple levels of repertoire organization, AbSolution enables systematic dissection of selection-driven changes in sequence-derived properties in adaptive immune repertoires.

## Discussion

Here we present AbSolution, an interactive and extensible R Shiny application for the systematic analysis and exploration of sequence-derived features in adaptive immune receptor repertoires. AbSolution uniquely integrates (i) large-scale sequence-derived feature computation, (ii) multiscale repertoire stratification, and (iii) interactive statistical feature selection within a single analysis environment. Its modular workflow and interlinked visualizations support accessible and dynamic analyses, enabling flexible hypothesis-driven and discovery-oriented exploration of immune repertoires. Together, these capabilities allow the exploration and analysis of clones and sequence features across experimental conditions, as demonstrated for both T-cell and B-cell repertoire datasets.

Our results show that immune selection leaves coherent, multiscale imprints on receptor sequence properties that cannot be captured by analyses focused on individual features, regions or single biological scales. Immune repertoires are highly heterogeneous, structured by clonal relationships, and analyses can be influenced by experimental design, sequencing depth, and gene usage biases. AbSolution, therefore, incorporates subsampling-based normalization strategies to mitigate these effects when comparing feature distributions between samples or experimental groups. These procedures allow more robust comparisons across samples while considering the underlying biological variation as we showed in our analysis of the B-cell repertoire. In addition, by integrating germline reconstruction with repertoire-level analyses, we show that these signatures reflect both inherited sequence constraints and SHM-driven adaptation.

In addition to the functionalities presented in the *Results* section, AbSolution also supports configurable filtering and normalization strategies, several sequence motif- and composition-based features, feature changes between repertoire and germline values for SHM-focused analysis. AbSolution also allows the automatic export of code, figures, and metadata required to reproduce an analysis and to facilitate transparent reporting (*manuscript submitted*).

Our current implementation employs univariate logistic regression with multiple testing correction to identify discriminative features between two groups. This approach prioritizes interpretability, scaling and robustness in high-dimensional settings but does not account for feature interactions or correlated covariates. Feature matrices, lists of significant features and metadata can be used for further modelling in external statistical or machine-learning frameworks (Akbar et al., 2021; Zaslavsky et al., 2025; Zhao et al., 2023). This design balances accessibility and ease-of-use within AbSolution with analytical flexibility provided by software such as R/RStudio. Future extensions incorporating multivariable modelling, methods for handling correlated features, and interaction analysis would reduce reliance on external tools and support more transparent, reproducible end-to-end workflows within AbSolution.

Several feature-related extensions could further expand the analytical scope of AbSolution. These include the incorporation of region-specific subsegments such as the DE-loop (Kelow et al., 2020) or the centre of the CDR3 (Egorov et al., 2018); additional structural descriptors (Cruciani et al., 2004; Postovskaya et al., 2024; Sheng et al., 2017; van Westen et al., 2013), and expanded k-mer representation (Greiff et al., 2020) that capture higher-order sequence patterns. However, integrating such features also introduces challenges, including increased computational complexity and execution time, redundancy between correlated descriptors, and the need for better visualization strategies to facilitate operating with high-dimensional outputs. Addressing these challenges will require careful prioritization of descriptors and modular expansion of the feature-calculation pipeline.

Beyond sequence-level analyses, future versions of AbSolution could support integration with complementary data modalities, including (sub)clonal analyses, paired-chain repertoires and single-cell transcriptomics. Computing and visualizing B-cell clonal and subclonal diversity metrics and phylogenies would enable systematic in-depth analysis of sequence evolution and feature trajectories during affinity maturation (Gupta et al., 2017). Extensions to paired-chain analysis would enable investigation of inter-chain relationships that influence antigen recognition and affinity-based selection (Jaffe et al., 2022; Seidler et al., 2025). Gene expression signatures have been shown to predict antibody affinity (Chirichella et al., 2026) and their usage could therefore provide an additional layer of information to interpret clonal expansion and the identification of key sequence-based features. For example, gene expression measurements could be incorporated as sample- or clone-associated metadata and analysed together with repertoire features, enabling correlation-based analyses or comparisons between high-and low-expression groups. However, integrating these modalities would require algorithms and workflows that support multiscale analysis while preserving linked selections, coordinated views, and intuitive exploration across data modalities.

In summary, AbSolution provides a versatile and reproducible platform for multiscale exploration of immune receptor repertoires. By combining flexible stratification of repertoires with accessible systematic characterization of sequence-derived properties, it advances our capacity to explore and interpret complex repertoire datasets and to generate mechanistic hypotheses regarding adaptive immune responses in health and disease.

## Materials and methods

### Dataset

Pre-processed sequences in AIRR-seq format from the TCR sequencing dataset PRJNA685965 (Pollastro et al., 2020) were obtained from the authors of the paper. Incomplete TCR sequences were aligned to germline V(D)J segments using IMGT/HighV-QUEST (Brochet et al., 2008), and missing variable-region portions were reconstructed from the germline references via a custom script outside AbSolution and archived together with the analysis code [https://doi.org/10.5281/zenodo.20121327]. Samples from BCR sequencing dataset PRJNA814462 (Anang et al., 2023) were pre-processed into AIRR-seq format using the Immcantation framework tools pRESTO v0.7.1 (Vander Heiden et al., 2014), Change-O v1.3.0, *alakazam* v1.2.1 and SHazaM v1.1.2 (Gupta et al., 2015).

### Workflow

A summary of steps and internal workflow for AbSolution is provided in *Figure 3*. Here we explain the main aspects and refer to the manual (*Supplementary file 1*) for more detailed information and functionality.

**Figure 3.**
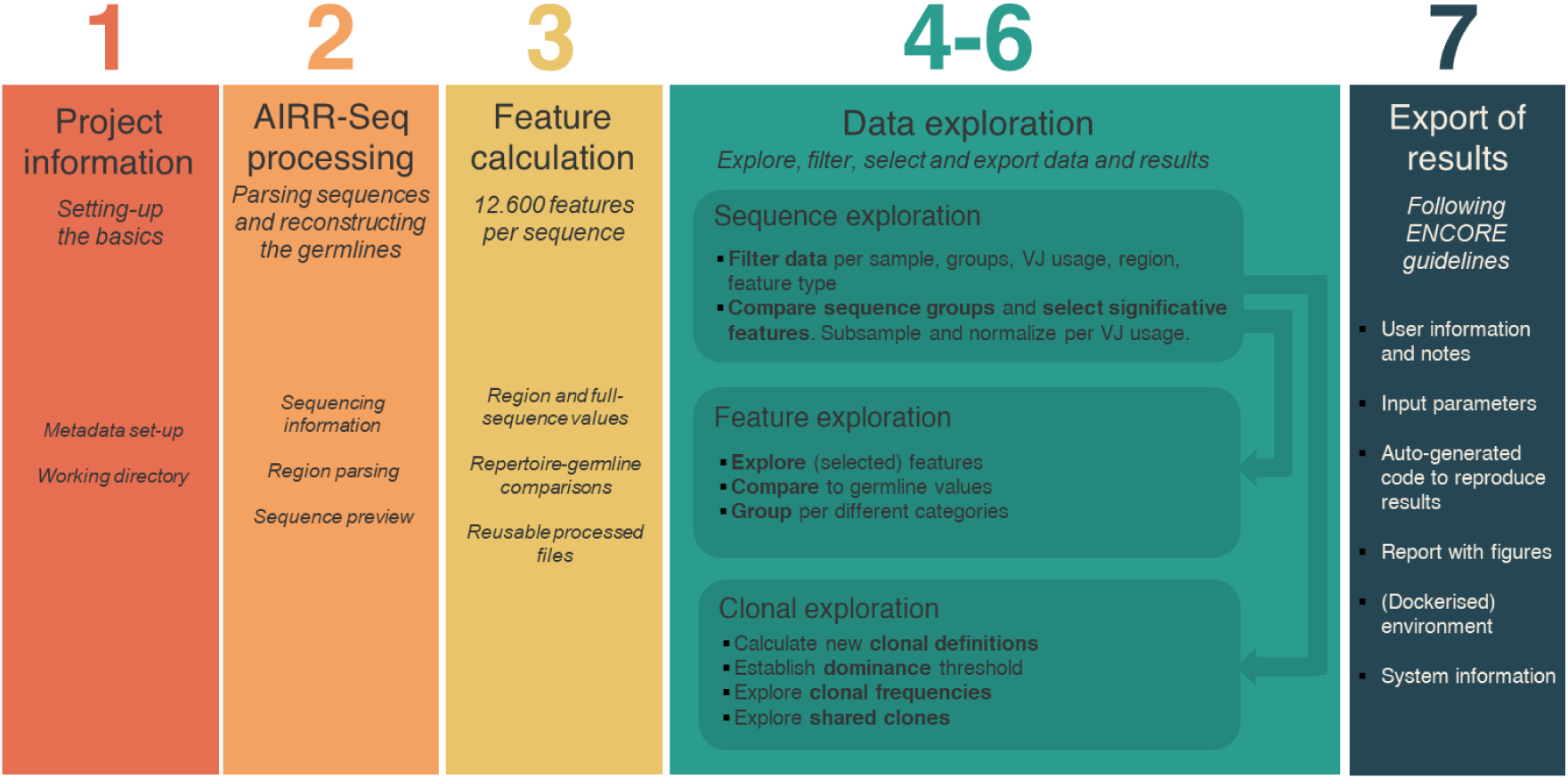
The AbSolution workflow. T-cell or B-cell AIRR-seq datasets are imported together with basic project and sequence metadata. Repertoire sequences are parsed and germline sequences are reconstructed using user-provided information on partial FWR1/4 regions, C-region presence, and D-gene alignment usage. An extensive set of sequence-derived features are calculated, many normalized or expressed relative to their germline values. The workflow is designed for reproducible reuse across analyses. When previously processed sequence information and feature tables are available, users can reload these files directly, allowing AIRR-seq processing and feature calculation (steps 2–3) to be omitted. The data-exploration interface comprises three interconnected sections: (1) ‘Sequence exploration’, for sequence and feature filtering followed by group comparisons, (2) ‘Feature exploration’, for visualizing features by group and comparison against germline values, and (3) ‘Clonal exploration’, which supports customizable clone definitions, dominance grouping by customized thresholds, and shared-clone analyses. Filters applied in the ‘Sequence exploration’ view automatically propagate to the Feature and Clonal sections. All user inputs, parameters and notes are automatically captured, enabling one-click export of reproducible R code snippets, publication-quality figures and a comprehensive HTML report that includes system information and package versioning.

### Step 1. Project information

BCR or TCR datasets are loaded in AIRR-seq standard format (Rubelt et al., 2017; Vander Heiden et al., 2018) and associated project metadata information (e.g., filename, sample type, subject identifier, group, and subgroup) is provided in tab-delimited format. To get acquainted with AbSolution, a BCR dataset from the *alakazam* package (Gupta et al., 2015) is included within AbSolution.

### Step 2. AIRR-seq processing

For each sequence, AbSolution reconstructs its corresponding germline sequence by mapping the observed FWR and CDR boundaries onto the reference germline, removing alignment gaps, and resolving ambiguous nucleotides in the CDR3 by replacing them with the corresponding bases from the repertoire sequence (optionally spanning the entire D region). The reconstruction can optionally span the entire D region, or instead retain the sequenced segment. Sequences carrying non-canonical nucleotides (including IUPAC ambiguity codes), or lacking complete FWR or CDR regions of sufficient length for reliable translation, are excluded. Sequences with stop codons in the ORF are likewise excluded. Because each repertoire sequence is analysed together with its reconstructed germline counterpart as a linked pair, removal of one automatically entails removal of the corresponding partner. To ensure compatibility with specific primer designs, AbSolution can be configured to retain sequences with partial FWR1 and/or FWR4 regions. If a C segment is present, it is removed during preprocessing.

### Step 3. Feature calculation

For each individual region (FWR1-4 and CDR1-3) and for the full V region of the TCR/BCR repertoire sequence and the reconstructed germline, the following features are computed:

i. *Physicochemical amino acid properties*. These include charge (Bjellqvist et al., 1993) at blood pH, normalized charge, bulkiness (Zimmerman et al., 1968), hydrophobicity index (Kyte & Doolittle, 1982), aliphatic index (Ikai, 1980), average polarity (Grantham, 1974), isoelectric point (Bjellqvist et al., 1993), Boman index (Boman, 2003), hydrophobic moment (Eisenberg et al., 1982), molecular weight (Wilkins et al., 1999), instability index (Guruprasad et al., 1990). We calculate these 11 properties using the *alakazam* (Gupta et al., 2015) and *Peptides* (Osorio et al., 2015) R packages. The value of a given property is obtained by summing the corresponding per-residue property values over all amino acids in the sequence and normalizing by its sequence length. These properties are additionally reported as differences relative to the corresponding reconstructed germline values.
ii. *SHM-related nucleotide and amino acid properties.* These include number and length of insertions and deletions (indels), number of replacement (R), number of silent (S) and total mutations, R/S ratio representing selection pressure (Shlomchik et al., 1987), number of observed transitions (exchange of purine to purine or pyrimidine to pyrimidine), transversions (exchange of purine and pyrimidine) and ratio of transitions and transversions (Holmquist, 1983) and number of specific nucleotide (NT)-to-NT (e.g., A to T), codon-to-codon (e.g., AAA to ATA, only for the complete variable sequence), amino acid (AA)-to-AA (e.g., proline to serine), and AA-class-to-AA-class (e.g., basic to acidic) exchanges to explore biased substitutions (Sheng et al., 2017). For each sequence, the total number of events in each category is computed, yielding 4,507 SHM-related features. With the exception of indel length, all properties are calculated relative to the corresponding reconstructed germline sequence. For each of these properties, values are additionally expressed in a length-normalized form, obtained by dividing by the corresponding sequence length.
iii. *Sequence motifs and composition*. This includes motifs (N-glycosylation sites (van de Bovenkamp et al., 2018) and SHM hot- and cold-spots (Yaari et al., 2013)), NT and AA sequence length (Sankar et al., 2020) and composition (NT, codon and AA usage, and AA classes). In total we compute 110 properties. For each of these properties, values are additionally expressed in a length-normalized form, obtained by dividing by the corresponding sequence length. These properties are additionally reported as differences relative to the corresponding reconstructed germline values.
iv. *Protein averages of amino acid descriptor sets* (van Westen et al., 2013). These include Kidera-Factors (Kidera et al., 1985), MS-WHIM scores (Zaliani & Gancia, 1999), FASGAI vectors (Liang et al., 2008), BLOSUM indices obtained by combining the VARIMAX analysis of AA physicochemical properties and the BLOSUM62 substitution matrix (Georgiev, 2009), VHSE-scales (Mei et al., 2005), Cruciani properties (Cruciani et al., 2004), ST-scales (Yang et al., 2010), T-scales (Tian et al., 2007) and Z-scales (Sandberg et al., 1998). These average values for each of the 58 properties are computed for each sequence using the *Peptides* R package (Osorio et al., 2015). These properties are additionally reported as differences relative to the corresponding reconstructed germline values.

A summary of the property classes can be found in *Table 3*. A summary of the 12.600 features calculated for these properties can be found in *Supplementary Table 1*. The features measured as differences between feature values in the repertoire and in the corresponding germline sequence enable quantitative assessment of sequence evolution of BCRs during the GC reaction. Many of these features are correlated (e.g., net negative charge correlates with the number of negatively charged residues). Nevertheless, all are retained for completeness. AbSolution also allows optional inclusion or exclusion of selected features during its downstream analyses.

**Table 3.** Overview of calculated features grouped per category. . Features are computed for both individual regions and the entire variable sequence. Composition, SHM-related and motif properties are additionally calculated as values normalized by sequence length. Physicochemical amino-acid and SHM-related nucleotide and amino-acid properties are also reported as differences relative to their germline values.

| Category | Description | Features |
| --- | --- | --- |
| **Physicochemical amino acid properties** | Properties related to amino acid characteristics are averaged over the sequence. | Charge, bulkiness, hydrophobicity index, aliphatic index, average polarity, isoelectric point, Boman index, hydrophobic moment, molecular weight, instability index. |
| **SHM-related nucleotide and amino acid properties** | Metrics related to the effect of somatic hypermutation (SHM). For BCRs only. | Number of insertions/deletions, replacement/silent/total mutations, R/S ratio (selection pressure), specific transitions/transversions (A↔T, C↔G), codon and amino acid transitions. |
| **Sequence motifs and composition** | Motif and sequence properties. | N-glycosylation sites, SHM hot/cold-spots, nucleotide and amino acid sequence lengths, NT/codon/AA usage, amino acid classes. |
| **Amino acid descriptor sets** | Physicochemical and structural vector sets for amino acids. | Kidera factors, MS-WHIM scores, FASGAI scales, BLOSUM scores, VHSE scales, Cruciani properties, ST-scales, T-scales, Z-scales. |

Additionally, all non-quantitative sequence metadata -including sequence identifiers, sample and patient information, experimental group, V(D)J gene usage, reading frame used, nucleotide and amino-acid sequences by region, and hotspot/cold-spot and N-glycosylation motif locations-is stored in the tab-delimited *Sequence information* table.

### Steps 4-6. Sequence, feature and clonal data exploration

In the *Sequence Exploration* tab, data may be interactively filtered by sequence type, sample, chain type, V-J usage, number of mutations, sequence region(s), and sequence-derived features. Feature variables with no variance are automatically discarded. Features can be filtered based on statistical significance using a feature-wise logistic regression module based on file-backed matrices (Prive et al., 2018) for specific binary comparisons (e.g., dominant vs non-dominant) (*Eq. 1*). Supplied sequence class labels can be employed as categorical annotations. Significance is assessed using a p-value threshold (default p ≤ 0.05), with optional multiple-testing correction using the Holm (Holm, 1979), Benjamini-Hochberg (Benjamini & Hochberg, 1995) or Benjamini-Yekutieli (Benjamini & Yekutieli, 2005) methods.

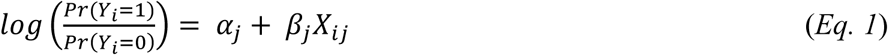

where

Y_i_ ∈ {0, 1} is the binary outcome for sequence i,

α_j_ is the intercept,

X_ij_ is the value of feature j for sequence i,

β_j_ is the effect size (slope) for sequence j

Multiple sampling schemes to enable robust comparisons are supported: (i) sequences can be subsampled at the clone level to remove clonal bias (Weinstein et al., 2013), (ii) sequences with identical VJ gene usage can be subsampled to avoid VJ bias (Marcou et al., 2018) and (iii) sequences can be subsampled across experimental groups to prevent overrepresentation of samples with higher sequencing depth. Each subsampling run is performed as a single random draw using the specified random seed, enabling reproducible subset selection. These subsampling-based normalization strategies are integrated within a decision framework for defining comparison settings *(Figure 4*). The framework enables standardized definition of comparison settings and ensures consistent application of normalization and subsampling procedures across analyses.

**Figure 4.**
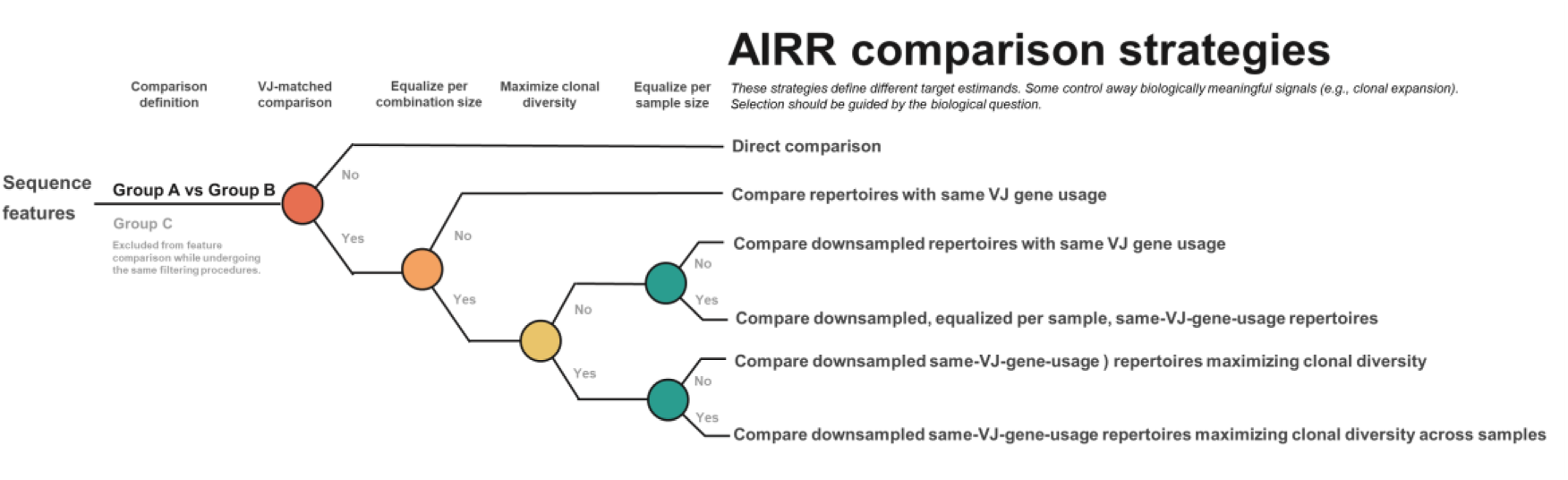
Workflow for defining AIRR repertoire comparison strategies in AbSolution. This diagram summarizes the decision framework used in AbSolution to define repertoire comparison strategies. Starting from an initial group-level comparison (Group A vs Group B), analyses proceed through successive filtering and subsampling-based normalization steps based on VJ gene matching, equalization of VJ combination size, maximization of clonal diversity, and balancing of sample contributions. At each decision point, progressively stricter controls reduce effects arising from differences in sequencing depth, VJ composition, clonal dominance, and sample-specific overrepresentation. Depending on the selected path, comparisons range from direct, unfiltered analyses to downsampled, VJ-matched repertoires with controlled clonal and sample representation. Sequences in group C are excluded from the group-level feature comparison while undergoing the same filtering procedures.

Different comparison strategies within the framework correspond to progressively stricter normalization settings and address distinct analytical questions. Direct comparisons preserve the full repertoire signal and allow detection of overall group differences, including changes in V/J gene usage and clonal expansion, but may be confounded by differences in sequencing depth, V/J composition, clonal dominance, or uneven sample representation. Restricting analyses to sequences with the same VJ gene usage controls for V/J composition and enables comparison within a matched VJ background, although other sources of bias such as sequencing depth or clonal expansion may remain. Downsampling of sequences with the same VJ gene usage further controls for differences in sequence abundance and sequencing depth between groups. Additional normalization steps can balance sample contributions, preventing individual samples from dominating group-level signals, or prioritize clonal diversity, reducing the influence of highly expanded clones and emphasizing broader repertoire trends. The most stringent strategy combines these controls, simultaneously accounting for V/J composition, sequencing depth, clonal dominance, and sample representation. While increasingly restrictive normalization improves robustness of comparisons, it also reduces the number of sequences available for analysis and excludes repertoire features outside the shared VJ space.

### Selection and Exploration sets

Two parallel working sets are maintained: the *Selection set*, which serves as the reference for downstream analyses, and the *Exploration set*, which may be subjected to alternative filtering parameters. Each set is visualized via principal component analysis (PCA), with an optional UMAP projection (McInnes et al., 2018) when data size permits in-memory computation. To facilitate interpretation, AbSolution implements a coloring algorithm that assigns similar colors to sequences sharing the same V, D, or J gene families, or identical VJ/VDJ family combinations. For each of the two sets, AbSolution reports sample metadata, V(D)J gene usage, retained features, and statistically significant Pearson correlations among these features with correlation coefficients (r) ≥ 0.8.

The interface displays two dimensionality-reduction plots side by side: a *Selection* panel showing the sequences included in the current analysis and an *Exploration* panel that allows users to test alternative filtering parameters and selections without altering the primary analysis. Selections made in the dimensionality-reduction plots of either the *Selection* or *Exploration set* are dynamically linked. Selecting sequences in one projection automatically highlights the corresponding repertoire and reconstructed germline sequences in the parallel set, updates the associated metadata, feature, and correlation tables, and enables export of the selected entries. This coordinated, linked visualization supports iterative, feature-driven exploration and direct comparison between reference and alternatively filtered sequence sets.

For the sequences in the *Selection set*, the distribution of each retained feature is summarized using violin plot representations, with optional stratification by experimental groupings. Reconstructed germline feature values are included as reference baselines, enabling direct comparison between repertoire and germline-derived measurements and facilitating identification of shifts in features. Statistical significance between the repertoire sequences and reconstructed germline sequence is assessed using two-sample t-tests, and effect size is measured by Cohen’s d.

Sequences in the *Selection set* are assigned to clonal groups based either on provided clone identifiers or on *de novo* clustering defined by shared V-J gene usage and identical CDR3 sequences at the nucleotide or amino-acid level, performed either per sample or across the full dataset for shared clones. Clonal abundance relative frequencies are calculated per sample, and depicted as split-violin plots for each sample. Dominant clones can be defined using a user-specified relative frequency threshold, while shared clonotypes across samples are identified and summarized using set-intersection UpSet plots (Lex et al., 2014). Interaction with the UpSet plot generates comparative frequency plots for the corresponding samples, distinguishing clones shared across all samples, shared between pairs of samples, and sample-specific or unique clones. Clonal grouping can be interactively defined from any of these visualizations to retrieve the corresponding repertoire and reconstructed germline sequences, enabling clone-level comparison of sequence features and subsequent feature-based stratification of clonal populations.

### Step 7. Export of results

After a complete analysis, data, code, software environment, system information, a report and results generated can be exported according to the ENCORE standardized file system structure, which was developed to enhance reproducibility (van Kampen et al., 2024).

### Software description

AbSolution is an R (R Core Team, 2021) package publicly available in CRAN (https://cran.r-project.org/package=AbSolution) for installation. It is a Shiny (Chang et al., 2024) application that supports data exploration and analysis of larger-than-RAM datasets through the use of memory-mapped matrices (R package *bigstatsr (Prive et al., 2018)*) and linear-scaling algorithms and techniques where possible. AbSolution follows the ENCORE (van Kampen et al., 2024) workflow for reproducibility. Its ENCORE development folder can be found at https://github.com/EDS-Bioinformatics-Laboratory/AbSolution.

## Acknowledgements

This work is supported by COSMIC (https://www.cosmic-h2020.eu), which has received funding from the European Union’s Horizon 2020 research and innovation programme under the Marie Skłodowska-Curie grant agreement No 765158. Dataset PRJNA685965 was generated by the ABIRISK consortium (Anti-Biopharmaceutical Immunization: Prediction and analysis of clinical relevance to minimize the risk; www.abirisk.eu) in collaboration with Prof. dr. Bernard Maillere, University Paris-Saclay, Department of Medicine and Health Technology, France. CT’s mobility was co-funded by the European Union under the Erasmus+ Traineeship scheme (2019-2020).

## Contributions

R.G.V., Av.K, H.H., J.G., D.A. and Nd.V defined the AbSolution framework. R.G.V. designed and led the implementation of AbSolution. C.T. contributed to the development of an early proof-of-concept. A.J. helped review and test the programme. S.P., D.A. and Nd.V. produced the datasets. Bv.S. processed the datasets. Sam Langton conducted a CODECHECK report. All authors contributed to the manuscript.

## Data availability

The datasets PRJNA685965 (Pollastro et al., 2020) and PRJNA814462 (Anang et al., 2023) are available from the NCBI BioProject repository.

## Code availability

Source code used for this work are available from GitHub [https://github.com/EDS-Bioinformatics-Laboratory/AbSolution] and CRAN (https://cran.r-project.org/package=AbSolution).

Code, raw, metadata and processed files of the analyses are available from Zenodo [https://doi.org/10.5281/zenodo.20121327].

## References

1. Akbar, R., Robert, P. A., Pavlovic, M., Jeliazkov, J. R., Snapkov, I., Slabodkin, A., Weber, C. R., Scheffer, L., Miho, E., Haff, I. H., Haug, D. T. T., Lund-Johansen, F., Safonova, Y., Sandve, G. K., & Greiff, V. (2021). A compact vocabulary of paratope-epitope interactions enables predictability of antibody-antigen binding. Cell Rep, 34(11), 108856. 10.1016/j.celrep.2021.108856

2. Alt, F. W., & Baltimore, D. (1982). Joining of immunoglobulin heavy chain gene segments: implications from a chromosome with evidence of three D-JH fusions. Proc Natl Acad Sci U S A, 79(13), 4118–4122. 10.1073/pnas.79.13.4118

3. Alt, F. W., Bothwell, A. L., Knapp, M., Siden, E., Mather, E., Koshland, M., & Baltimore, D. (1980). Synthesis of secreted and membrane-bound immunoglobulin mu heavy chains is directed by mRNAs that differ at their 3’ ends. Cell, 20(2), 293–301. 10.1016/0092-8674(80)90615-7

4. Anang, D. C., Walter, H. A. W., Lim, J., Niewold, I., van der Weele, L., Aronica, E., Eftimov, F., Raaphorst, J., van Schaik, B. D. C., van Kampen, A. H. C., van der Kooi, A. J., & de Vries, N. (2023). B-cell receptor profiling before and after IVIG monotherapy in newly diagnosed idiopathic inflammatory myopathies. Rheumatology (Oxford), 62(7), 2585–2593. 10.1093/rheumatology/keac602

5. Andrade, D. S., Terrematte, P., Renno-Costa, C., Zilberberg, A., & Efroni, S. (2023). GENTLE: a novel bioinformatics tool for generating features and building classifiers from T cell repertoire cancer data. BMC Bioinformatics, 24(1), 32. 10.1186/s12859-023-05155-w

6. Arnaout, R. A., Prak, E. T. L., Schwab, N., Rubelt, F., & Adaptive Immune Receptor Repertoire, C. (2021). The Future of Blood Testing Is the Immunome. Front Immunol, 12, 626793. 10.3389/fimmu.2021.626793

7. Bashford-Rogers, R. J. M., Bergamaschi, L., McKinney, E. F., Pombal, D. C., Mescia, F., Lee, J. C., Thomas, D. C., Flint, S. M., Kellam, P., Jayne, D. R. W., Lyons, P. A., & Smith, K. G. C. (2019). Analysis of the B cell receptor repertoire in six immune-mediated diseases. Nature, 574(7776), 122–126. 10.1038/s41586-019-1595-3

8. Benjamini, Y., & Hochberg, Y. (1995). Controlling the False Discovery Rate: A Practical and Powerful Approach to Multiple Testing. Journal of the Royal Statistical Society Series B: Statistical Methodology, 57(1), 289–300. 10.1111/j.2517-6161.1995.tb02031.x

9. Benjamini, Y., & Yekutieli, D. (2005). False Discovery Rate–Adjusted Multiple Confidence Intervals for Selected Parameters. Journal of the American Statistical Association, 100(469), 71–81. 10.1198/016214504000001907

10. Bjellqvist, B., Hughes, G. J., Pasquali, C., Paquet, N., Ravier, F., Sanchez, J. C., Frutiger, S., & Hochstrasser, D. (1993). The focusing positions of polypeptides in immobilized pH gradients can be predicted from their amino acid sequences. Electrophoresis, 14(10), 1023–1031. 10.1002/elps.11501401163

11. Boman, H. G. (2003). Antibacterial peptides: basic facts and emerging concepts. J Intern Med, 254(3), 197–215. 10.1046/j.1365-2796.2003.01228.x

12. Boyd, S. D., & Crowe, J. E., Jr. (2016). Deep sequencing and human antibody repertoire analysis. Curr Opin Immunol, 40, 103–109. 10.1016/j.coi.2016.03.008

13. Brenner, M. B., McLean, J., Dialynas, D. P., Strominger, J. L., Smith, J. A., Owen, F. L., Seidman, J. G., Ip, S., Rosen, F., & Krangel, M. S. (1986). Identification of a putative second T-cell receptor. Nature, 322(6075), 145–149. 10.1038/322145a0

14. Brochet, X., Lefranc, M. P., & Giudicelli, V. (2008). IMGT/V-QUEST: the highly customized and integrated system for IG and TR standardized V-J and V-D-J sequence analysis. Nucleic Acids Res, 36(Web Server issue), W503-508. 10.1093/nar/gkn316

15. Busch, D. H., & Pamer, E. G. (1999). T cell affinity maturation by selective expansion during infection. J Exp Med, 189(4), 701–710. 10.1084/jem.189.4.701

16. Chang, W., Cheng, J., Allaire, J. J., Sievert, C., Schloerke, B., Xie, Y., Allen, J., McPherson, J., Dipert, A., & Borges, B. (2024). shiny: Web Application Framework for R. In https://shiny.posit.co/

17. Chirichella, M., Ratcliff, M., Gu, S., Miragaia, R. J., Sammito, M., Cutano, V., Cohen, S., Angeletti, D., Romero-Ros, X., & Schofield, D. J. (2026). Integrated single-cell analyses of affinity-tested B cells enable the identification of a gene signature to predict antibody affinity. Cell Syst, 17(2), 101483. 10.1016/j.cels.2025.101483

18. Crescioli, S., Correa, I., Ng, J., Willsmore, Z. N., Laddach, R., Chenoweth, A., Chauhan, J., Di Meo, A., Stewart, A., Kalliolia, E., Alberts, E., Adams, R., Harris, R. J., Mele, S., Pellizzari, G., Black, A. B. M., Bax, H. J., Cheung, A., Nakamura, M., . . . Karagiannis, S. N. (2023). B cell profiles, antibody repertoire and reactivity reveal dysregulated responses with autoimmune features in melanoma. Nat Commun, 14(1), 3378. 10.1038/s41467-023-39042-y

19. Cruciani, G., Baroni, M., Carosati, E., Clementi, M., Valigi, R., & Clementi, S. (2004). Peptide studies by means of principal properties of amino acids derived from MIF descriptors. Journal of Chemometrics, 18(3-4), 146–155. 10.1002/cem.856

20. Desiderio, S. V., Yancopoulos, G. D., Paskind, M., Thomas, E., Boss, M. A., Landau, N., Alt, F. W., & Baltimore, D. (1984). Insertion of N regions into heavy-chain genes is correlated with expression of terminal deoxytransferase in B cells. Nature, 311(5988), 752–755. 10.1038/311752a0

21. Dong, Y., Pi, X., Bartels-Burgahn, F., Saltukoglu, D., Liang, Z., Yang, J., Alt, F. W., Reth, M., & Wu, H. (2022). Structural principles of B cell antigen receptor assembly. Nature, 612(7938), 156–161. 10.1038/s41586-022-05412-7

22. Dupic, T., Marcou, Q., Walczak, A. M., & Mora, T. (2019). Genesis of the alphabeta T-cell receptor. PLoS Comput Biol, 15(3), e1006874. 10.1371/journal.pcbi.1006874

23. Edelman, G. M., Cunningham, B. A., Gall, W. E., Gottlieb, P. D., Rutishauser, U., & Waxdal, M. J. (1969). The covalent structure of an entire gammaG immunoglobulin molecule. Proc Natl Acad Sci U S A, 63(1), 78–85. 10.1073/pnas.63.1.78

24. Edelman, G. M., & Poulik, M. D. (1961). Studies on structural units of the gamma-globulins. J Exp Med, 113(5), 861–884. 10.1084/jem.113.5.861

25. Egorov, E. S., Kasatskaya, S. A., Zubov, V. N., Izraelson, M., Nakonechnaya, T. O., Staroverov, D. B., Angius, A., Cucca, F., Mamedov, I. Z., Rosati, E., Franke, A., Shugay, M., Pogorelyy, M. V., Chudakov, D. M., & Britanova, O. V. (2018). The Changing Landscape of Naive T Cell Receptor Repertoire With Human Aging. Front Immunol, 9, 1618. 10.3389/fimmu.2018.01618

26. Eisenberg, D., Weiss, R. M., & Terwilliger, T. C. (1982). The helical hydrophobic moment: a measure of the amphiphilicity of a helix. Nature, 299(5881), 371–374. 10.1038/299371a0

27. Georgiev, A. G. (2009). Interpretable numerical descriptors of amino acid space. J Comput Biol, 16(5), 703–723. 10.1089/cmb.2008.0173

28. Ghraichy, M., von Niederhausern, V., Kovaltsuk, A., Galson, J. D., Deane, C. M., & Truck, J. (2021). Different B cell subpopulations show distinct patterns in their IgH repertoire metrics. Elife, 10. 10.7554/eLife.73111

29. Goldstein, L. D., Chen, Y. J., Wu, J., Chaudhuri, S., Hsiao, Y. C., Schneider, K., Hoi, K. H., Lin, Z., Guerrero, S., Jaiswal, B. S., Stinson, J., Antony, A., Pahuja, K. B., Seshasayee, D., Modrusan, Z., Hotzel, I., & Seshagiri, S. (2019). Massively parallel single-cell B-cell receptor sequencing enables rapid discovery of diverse antigen-reactive antibodies. Commun Biol, 2, 304. 10.1038/s42003-019-0551-y

30. Grantham, R. (1974). Amino acid difference formula to help explain protein evolution. Science, 185(4154), 862–864. 10.1126/science.185.4154.862

31. Greiff, V., Yaari, G., & Cowell, L. G. (2020). Mining adaptive immune receptor repertoires for biological and clinical information using machine learning. Current Opinion in Systems Biology, 24, 109–119. 10.1016/j.coisb.2020.10.010

32. Gupta, N. T., Adams, K. D., Briggs, A. W., Timberlake, S. C., Vigneault, F., & Kleinstein, S. H. (2017). Hierarchical Clustering Can Identify B Cell Clones with High Confidence in Ig Repertoire Sequencing Data. J Immunol, 198(6), 2489–2499. 10.4049/jimmunol.1601850

33. Gupta, N. T., Vander Heiden, J. A., Uduman, M., Gadala-Maria, D., Yaari, G., & Kleinstein, S. H. (2015). Change-O: a toolkit for analyzing large-scale B cell immunoglobulin repertoire sequencing data. Bioinformatics, 31(20), 3356–3358. 10.1093/bioinformatics/btv359

34. Guruprasad, K., Reddy, B. V., & Pandit, M. W. (1990). Correlation between stability of a protein and its dipeptide composition: a novel approach for predicting in vivo stability of a protein from its primary sequence. Protein Eng, 4(2), 155–161. 10.1093/protein/4.2.155

35. Hochstenbach, F., & Brenner, M. B. (1989). T-cell receptor delta-chain can substitute for alpha to form a beta delta heterodimer. Nature, 340(6234), 562–565. 10.1038/340562a0

36. Holm, S. (1979). A Simple Sequentially Rejective Multiple Test Procedure. Scandinavian Journal of Statistics, 6(2), 6. 10.2307/4615733

37. Holmquist, R. (1983). Transitions and transversions in evolutionary descent: an approach to understanding. J Mol Evol, 19(2), 134–144. 10.1007/BF02300751

38. Hozumi, N., & Tonegawa, S. (1976). Evidence for somatic rearrangement of immunoglobulin genes coding for variable and constant regions. Proc Natl Acad Sci U S A, 73(10), 3628–3632. 10.1073/pnas.73.10.3628

39. Ikai, A. (1980). Thermostability and aliphatic index of globular proteins. J Biochem, 88(6), 1895–1898. 10.1093/OXFORDJOURNALS.JBCHEM.A133168

40. Jaffe, D. B., Shahi, P., Adams, B. A., Chrisman, A. M., Finnegan, P. M., Raman, N., Royall, A. E., Tsai, F., Vollbrecht, T., Reyes, D. S., Hepler, N. L., & McDonnell, W. J. (2022). Functional antibodies exhibit light chain coherence. Nature, 611(7935), 352–357. 10.1038/s41586-022-05371-z

41. Kasatskaya, S. A., Ladell, K., Egorov, E. S., Miners, K. L., Davydov, A. N., Metsger, M., Staroverov, D. B., Matveyshina, E. K., Shagina, I. A., Mamedov, I. Z., Izraelson, M., Shelyakin, P. V., Britanova, O. V., Price, D. A., & Chudakov, D. M. (2020). Functionally specialized human CD4(+) T-cell subsets express physicochemically distinct TCRs. Elife, 9. 10.7554/eLife.57063

42. Kawashima, S., & Kanehisa, M. (2000). AAindex: amino acid index database. Nucleic Acids Res, 28(1), 374. 10.1093/nar/28.1.374

43. Kelow, S. P., Adolf-Bryfogle, J., & Dunbrack, R. L. (2020). Hiding in plain sight: structure and sequence analysis reveals the importance of the antibody DE loop for antibody-antigen binding. MAbs, 12(1), 1840005. 10.1080/19420862.2020.1840005

44. Kidera, A., Konishi, Y., Oka, M., Ooi, T., & Scheraga, H. A. (1985). Statistical analysis of the physical properties of the 20 naturally occurring amino acids. Journal of Protein Chemistry, 4(1), 23–55. 10.1007/bf01025492

45. Klarenbeek, P. L., de Hair, M. J., Doorenspleet, M. E., van Schaik, B. D., Esveldt, R. E., van de Sande, M. G., Cantaert, T., Gerlag, D. M., Baeten, D., van Kampen, A. H., Baas, F., Tak, P. P., & de Vries, N. (2012). Inflamed target tissue provides a specific niche for highly expanded T-cell clones in early human autoimmune disease. Ann Rheum Dis, 71(6), 1088–1093. 10.1136/annrheumdis-2011-200612

46. Klarenbeek, P. L., Tak, P. P., van Schaik, B. D., Zwinderman, A. H., Jakobs, M. E., Zhang, Z., van Kampen, A. H., van Lier, R. A., Baas, F., & de Vries, N. (2010). Human T-cell memory consists mainly of unexpanded clones. Immunol Lett, 133(1), 42–48. 10.1016/j.imlet.2010.06.011

47. Kreslavsky, T., Gleimer, M., Garbe, A. I., & von Boehmer, H. (2010). alphabeta versus gammadelta fate choice: counting the T-cell lineages at the branch point. Immunol Rev, 238(1), 169–181. 10.1111/j.1600-065X.2010.00947.x

48. Kyte, J., & Doolittle, R. F. (1982). A simple method for displaying the hydropathic character of a protein. J Mol Biol, 157(1), 105–132. 10.1016/0022-2836(82)90515-0

49. Lafaille, J. J., DeCloux, A., Bonneville, M., Takagaki, Y., & Tonegawa, S. (1989). Junctional sequences of T cell receptor gamma delta genes: implications for gamma delta T cell lineages and for a novel intermediate of V-(D)-J joining. Cell, 59(5), 859–870. 10.1016/0092-8674(89)90609-0

50. Laffy, J. M. J., Dodev, T., Macpherson, J. A., Townsend, C., Lu, H. C., Dunn-Walters, D., & Fraternali, F. (2017). Promiscuous antibodies characterised by their physico-chemical properties: From sequence to structure and back. Prog Biophys Mol Biol, 128, 47–56. 10.1016/j.pbiomolbio.2016.09.002

51. Lefranc, M. P., Pommie, C., Ruiz, M., Giudicelli, V., Foulquier, E., Truong, L., Thouvenin-Contet, V., & Lefranc, G. (2003). IMGT unique numbering for immunoglobulin and T cell receptor variable domains and Ig superfamily V-like domains. Dev Comp Immunol, 27(1), 55–77. http://www.ncbi.nlm.nih.gov/pubmed/12477501

52. Lex, A., Gehlenborg, N., Strobelt, H., Vuillemot, R., & Pfister, H. (2014). UpSet: Visualization of Intersecting Sets. IEEE Trans Vis Comput Graph, 20(12), 1983–1992. 10.1109/TVCG.2014.2346248

53. Liang, G., Chen, G., Niu, W., & Li, Z. (2008). Factor analysis scales of generalized amino acid information as applied in predicting interactions between the human amphiphysin-1 SH3 domains and their peptide ligands. Chem Biol Drug Des, 71(4), 345–351. 10.1111/j.1747-0285.2008.00641.x

54. Liu, H., Pan, W., Tang, C., Tang, Y., Wu, H., Yoshimura, A., Deng, Y., He, N., & Li, S. (2021). The methods and advances of adaptive immune receptors repertoire sequencing. Theranostics, 11(18), 8945–8963. 10.7150/thno.61390

55. Marcou, Q., Mora, T., & Walczak, A. M. (2018). High-throughput immune repertoire analysis with IGoR. Nat Commun, 9(1), 561. 10.1038/s41467-018-02832-w

56. Margreitter, C., Lu, H. C., Townsend, C., Stewart, A., Dunn-Walters, D. K., & Fraternali, F. (2018). BRepertoire: a user-friendly web server for analysing antibody repertoire data. Nucleic Acids Res, 46(W1), W264–W270. 10.1093/nar/gky276

57. McInnes, L., Healy, J., Saul, N., & Großberger, L. (2018). UMAP: Uniform Manifold Approximation and Projection. Journal of Open Source Software, 3(29). 10.21105/joss.00861

58. Mei, H., Liao, Z. H., Zhou, Y., & Li, S. Z. (2005). A new set of amino acid descriptors and its application in peptide QSARs. Biopolymers, 80(6), 775–786. 10.1002/bip.20296

59. Michielin, O., Luescher, I., & Karplus, M. (2000). Modeling of the TCR-MHC-peptide complex. J Mol Biol, 300(5), 1205–1235. 10.1006/jmbi.2000.3788

60. Nikolich-Zugich, J., Slifka, M. K., & Messaoudi, I. (2004). The many important facets of T-cell repertoire diversity. Nat Rev Immunol, 4(2), 123–132. 10.1038/nri1292

61. Osorio, D., Rondón-Villarreal, P., & Torres, R. (2015). Peptides: A Package for Data Mining of Antimicrobial Peptides. The R Journal, 7(1). 10.32614/rj-2015-001

62. Peled, J. U., Kuang, F. L., Iglesias-Ussel, M. D., Roa, S., Kalis, S. L., Goodman, M. F., & Scharff, M. D. (2008). The biochemistry of somatic hypermutation. Annu Rev Immunol, 26, 481–511. 10.1146/annurev.immunol.26.021607.090236

63. Pollastro, S., de Bourayne, M., Balzaretti, G., Jongejan, A., van Schaik, B. D. C., Niewold, I. T. G., van Kampen, A. H. C., Maillere, B., & de Vries, N. (2020). Characterization and Monitoring of Antigen-Responsive T Cell Clones Using T Cell Receptor Gene Expression Analysis. Front Immunol, 11, 609624. 10.3389/fimmu.2020.609624

64. Pollastro, S., Musters, A., Balzaretti, G., Niewold, I., van Schaik, B., Hassler, S., Verhoef, C. M., Pallardy, M., van Kampen, A., Mariette, X., de Vries, N., & Consortium, A. (2024). Sensitive B-cell receptor repertoire analysis shows repopulation correlates with clinical response to rituximab in rheumatoid arthritis. Arthritis Res Ther, 26(1), 70. 10.1186/s13075-024-03297-7

65. Porter, R. R. (1959). The hydrolysis of rabbit y-globulin and antibodies with crystalline papain. Biochem J, 73(1), 119–126. 10.1042/bj0730119

66. Postovskaya, A., Vercauteren, K., Meysman, P., & Laukens, K. (2024). tcrBLOSUM: an amino acid substitution matrix for sensitive alignment of distant epitope-specific TCRs. Brief Bioinform, 26(1). 10.1093/bib/bbae602

67. Prive, F., Aschard, H., Ziyatdinov, A., & Blum, M. G. B. (2018). Efficient analysis of large-scale genome-wide data with two R packages: bigstatsr and bigsnpr. Bioinformatics, 34(16), 2781–2787. 10.1093/bioinformatics/bty185

68. R Core Team. (2021). R: A language and environment for statistical computing. R Foundation for Statistical Computing. In https://www.R-project.org/

69. Raphael, I., Xiong, Z., Sneiderman, C. T., Raphael, R. A., Mash, M., Schwegman, L., Jackson, S. A., O’Brien, C., Anderson, K. J., Sever, R. E., Hendrikse, L. D., Vincze, S. R., Diaz, A., Felker, J., Nazarian, J., Nechemia-Arbely, Y., Hu, B., Kammula, U. S., Agnihotri, S., . . . Kohanbash, G. (2025). The T cell receptor landscape of childhood brain tumors. Sci Transl Med, 17(790), eadp0675. 10.1126/scitranslmed.adp0675

70. Rees, A. R. (2020). Understanding the human antibody repertoire. MAbs, 12(1), 1729683. 10.1080/19420862.2020.1729683

71. Rosenberg, A. M., Ayres, C. M., Medina-Cucurella, A. V., Whitehead, T. A., & Baker, B. M. (2024). Enhanced T cell receptor specificity through framework engineering. Front Immunol, 15, 1345368. 10.3389/fimmu.2024.1345368

72. Roskin, K. M., Jackson, K. J. L., Lee, J. Y., Hoh, R. A., Joshi, S. A., Hwang, K. K., Bonsignori, M., Pedroza-Pacheco, I., Liao, H. X., Moody, M. A., Fire, A. Z., Borrow, P., Haynes, B. F., & Boyd, S. D. (2020). Aberrant B cell repertoire selection associated with HIV neutralizing antibody breadth. Nat Immunol, 21(2), 199–209. 10.1038/s41590-019-0581-0

73. Rubelt, F., Busse, C. E., Bukhari, S. A. C., Burckert, J. P., Mariotti-Ferrandiz, E., Cowell, L. G., Watson, C. T., Marthandan, N., Faison, W. J., Hershberg, U., Laserson, U., Corrie, B. D., Davis, M. M., Peters, B., Lefranc, M. P., Scott, J. K., Breden, F., Community, A., Luning Prak, E. T., & Kleinstein, S. H. (2017). Adaptive Immune Receptor Repertoire Community recommendations for sharing immune-repertoire sequencing data. Nat Immunol, 18(12), 1274–1278. 10.1038/ni.3873

74. Saito, H., Kranz, D. M., Takagaki, Y., Hayday, A. C., Eisen, H. N., & Tonegawa, S. (1984). Complete primary structure of a heterodimeric T-cell receptor deduced from cDNA sequences. Nature, 309(5971), 757–762. 10.1038/309757a0

75. Sandberg, M., Eriksson, L., Jonsson, J., Sjostrom, M., & Wold, S. (1998). New chemical descriptors relevant for the design of biologically active peptides. A multivariate characterization of 87 amino acids. J Med Chem, 41(14), 2481–2491. 10.1021/jm9700575

76. Sankar, K., Hoi, K. H., & Hötzel, I. (2020). Dynamics of heavy chain junctional length biases in antibody repertoires. Communications Biology, 3(1). 10.1038/s42003-020-0931-3

77. Schroeder, H. W., Jr., & Cavacini, L. (2010). Structure and function of immunoglobulins. J Allergy Clin Immunol, *125*(2 Suppl 2), S41-52. 10.1016/j.jaci.2009.09.046

78. Seidler, C. A., Spanke, V. A., Gamper, J., Bujotzek, A., Georges, G., & Liedl, K. R. (2025). Data-driven analyses of human antibody variable domain germlines: pairings, sequences and structural features. MAbs, 17(1), 2507950. 10.1080/19420862.2025.2507950

79. Sela-Culang, I., Kunik, V., & Ofran, Y. (2013). The structural basis of antibody-antigen recognition. Front Immunol, 4, 302. 10.3389/fimmu.2013.00302

80. Setliff, I., Shiakolas, A. R., Pilewski, K. A., Murji, A. A., Mapengo, R. E., Janowska, K., Richardson, S., Oosthuysen, C., Raju, N., Ronsard, L., Kanekiyo, M., Qin, J. S., Kramer, K. J., Greenplate, A. R., McDonnell, W. J., Graham, B. S., Connors, M., Lingwood, D., Acharya, P., . . . Georgiev, I. S. (2019). High-Throughput Mapping of B Cell Receptor Sequences to Antigen Specificity. Cell, 179(7), 1636–1646 e1615. 10.1016/j.cell.2019.11.003

81. Shcherbinin, D. S., Karnaukhov, V. K., Zvyagin, I. V., Chudakov, D. M., & Shugay, M. (2023). Large-scale template-based structural modeling of T-cell receptors with known antigen specificity reveals complementarity features. Front Immunol, 14, 1224969. 10.3389/fimmu.2023.1224969

82. Sheng, Z., Schramm, C. A., Kong, R., Program, N. C. S., Mullikin, J. C., Mascola, J. R., Kwong, P. D., & Shapiro, L. (2017). Gene-Specific Substitution Profiles Describe the Types and Frequencies of Amino Acid Changes during Antibody Somatic Hypermutation. Front Immunol, 8, 537. 10.3389/fimmu.2017.00537

83. Shlomchik, M. J., Marshak-Rothstein, A., Wolfowicz, C. B., Rothstein, T. L., & Weigert, M. G. (1987). The role of clonal selection and somatic mutation in autoimmunity. Nature, 328(6133), 805–811. 10.1038/328805a0

84. Shugay, M., Bagaev, D. V., Turchaninova, M. A., Bolotin, D. A., Britanova, O. V., Putintseva, E. V., Pogorelyy, M. V., Nazarov, V. I., Zvyagin, I. V., Kirgizova, V. I., Kirgizov, K. I., Skorobogatova, E. V., & Chudakov, D. M. (2015). VDJtools: Unifying Post-analysis of T Cell Receptor Repertoires. PLoS Comput Biol, 11(11), e1004503. 10.1371/journal.pcbi.1004503

85. Sofou, E., Vlachonikola, E., Zaragoza-Infante, L., Bruggemann, M., Darzentas, N., Groenen, P., Hummel, M., Macintyre, E. A., Psomopoulos, F., Davi, F., Langerak, A. W., & Stamatopoulos, K. (2023). Clonotype definitions for immunogenetic studies: proposals from the EuroClonality NGS Working Group. Leukemia, 37(8), 1750–1752. 10.1038/s41375-023-01952-7

86. Taft, J. M., Weber, C. R., Gao, B., Ehling, R. A., Han, J., Frei, L., Metcalfe, S. W., Overath, M. D., Yermanos, A., Kelton, W., & Reddy, S. T. (2022). Deep mutational learning predicts ACE2 binding and antibody escape to combinatorial mutations in the SARS-CoV-2 receptor-binding domain. Cell, 185(21), 4008–4022 e4014. 10.1016/j.cell.2022.08.024

87. Tatikonda, R. R., Demerdash, O. N. A., & Smith, J. C. (2024). TCR-H: explainable machine learning prediction of T-cell receptor epitope binding on unseen datasets. Front Immunol, 15, 1426173. 10.3389/fimmu.2024.1426173

88. Tian, F., Zhou, P., & Li, Z. (2007). T-scale as a novel vector of topological descriptors for amino acids and its application in QSARs of peptides. Journal of Molecular Structure, 830(1-3), 106–115. 10.1016/j.molstruc.2006.07.004

89. Todeschini, R., & Consonni, V. (2000). Handbook of Molecular Descriptors. Wiley-VCH. 10.1002/9783527613106

90. Townsend, C. L., Laffy, J. M., Wu, Y. B., Silva O’Hare, J., Martin, V., Kipling, D., Fraternali, F., & Dunn-Walters, D. K. (2016). Significant Differences in Physicochemical Properties of Human Immunoglobulin Kappa and Lambda CDR3 Regions. Front Immunol, 7, 388. 10.3389/fimmu.2016.00388

91. Tsuchiya, Y., Namiuchi, Y., Wako, H., & Tsurui, H. (2018). A study of CDR3 loop dynamics reveals distinct mechanisms of peptide recognition by T-cell receptors exhibiting different levels of cross-reactivity. Immunology, 153(4), 466–478. 10.1111/imm.12849

92. van de Bovenkamp, F. S., Derksen, N. I. L., Ooijevaar-de Heer, P., van Schie, K. A., Kruithof, S., Berkowska, M. A., van der Schoot, C. E. H. I. J. van der Burg, M., Gils, A., Hafkenscheid, L., Toes, R. E. M., Rombouts, Y., Plomp, R., Wuhrer, M., van Ham, S. M., Vidarsson, G., & Rispens, T. (2018). Adaptive antibody diversification through N-linked glycosylation of the immunoglobulin variable region. Proc Natl Acad Sci U S A, 115(8), 1901–1906. 10.1073/pnas.1711720115

93. van Kampen, A. H. C., Mahamune, U., Jongejan, A., van Schaik, B. D. C., Balashova, D., Lashgari, D., Pras-Raves, M., Wever, E. J. M., Dane, A. D., Garcia-Valiente, R., & Moerland, P. D. (2024). ENCORE: a practical implementation to improve reproducibility and transparency of computational research. Nat Commun, 15(1), 8117. 10.1038/s41467-024-52446-8

94. van Westen, G. J., Swier, R. F., Wegner, J. K., Ijzerman, A. P., van Vlijmen, H. W., & Bender, A. (2013). Benchmarking of protein descriptor sets in proteochemometric modeling (part 1): comparative study of 13 amino acid descriptor sets. J Cheminform, 5(1), 41. 10.1186/1758-2946-5-41

95. Vander Heiden, J. A., Marquez, S., Marthandan, N., Bukhari, S. A. C., Busse, C. E., Corrie, B., Hershberg, U., Kleinstein, S. H., Matsen Iv, F. A., Ralph, D. K., Rosenfeld, A. M., Schramm, C. A., Community, A., Christley, S., & Laserson, U. (2018). AIRR Community Standardized Representations for Annotated Immune Repertoires. Front Immunol, 9, 2206. 10.3389/fimmu.2018.02206

96. Vander Heiden, J. A., Yaari, G., Uduman, M., Stern, J. N., O’Connor, K. C., Hafler, D. A., Vigneault, F., & Kleinstein, S. H. (2014). pRESTO: a toolkit for processing high-throughput sequencing raw reads of lymphocyte receptor repertoires. Bioinformatics, 30(13), 1930–1932. 10.1093/bioinformatics/btu138

97. Waltari, E., Nafees, S., McCutcheon, K. M., Wong, J., & Pak, J. E. (2022). AIRRscape: An interactive tool for exploring B-cell receptor repertoires and antibody responses. PLoS Comput Biol, 18(9), e1010052. 10.1371/journal.pcbi.1010052

98. Weill, J. C., & Reynaud, C. A. (2020). IgM memory B cells: specific effectors of innate-like and adaptive responses. Curr Opin Immunol, 63, 1–6. 10.1016/j.coi.2019.09.003

99. Weinstein, J. A., Zeng, X., Chien, Y. H., & Quake, S. R. (2013). Correlation of gene expression and genome mutation in single B-cells. PLoS One, 8(6), e67624. 10.1371/journal.pone.0067624

100. Wilkins, M. R., Gasteiger, E., Bairoch, A., Sanchez, J. C., Williams, K. L., Appel, R. D., & Hochstrasser, D. F. (1999). Protein identification and analysis tools in the ExPASy server. Methods Mol Biol, 112, 531–552. 10.1385/1-59259-584-7:531

101. Wu, T. T., & Kabat, E. A. (1970). An analysis of the sequences of the variable regions of Bence Jones proteins and myeloma light chains and their implications for antibody complementarity. J Exp Med, 132(2), 211–250. https://www.ncbi.nlm.nih.gov/pubmed/5508247 http://jem.rupress.org/content/jem/132/2/211.full.pdf

102. Yaari, G., Vander Heiden, J. A., Uduman, M., Gadala-Maria, D., Gupta, N., Stern, J. N., O’Connor, K. C., Hafler, D. A., Laserson, U., Vigneault, F., & Kleinstein, S. H. (2013). Models of somatic hypermutation targeting and substitution based on synonymous mutations from high-throughput immunoglobulin sequencing data. Front Immunol, 4, 358. 10.3389/fimmu.2013.00358

103. Yancopoulos, G. D., Desiderio, S. V., Paskind, M., Kearney, J. F., Baltimore, D., & Alt, F. W. (1984). Preferential utilization of the most JH-proximal VH gene segments in pre-B-cell lines. Nature, 311(5988), 727–733. 10.1038/311727a0

104. Yang, L., Shu, M., Ma, K., Mei, H., Jiang, Y., & Li, Z. (2010). ST-scale as a novel amino acid descriptor and its application in QSAM of peptides and analogues. Amino Acids, 38(3), 805–816. 10.1007/s00726-009-0287-y

105. Zaliani, A., & Gancia, E. (1999). MS-WHIM Scores for Amino Acids: A New 3D-Description for Peptide QSAR and QSPR Studies. Journal of Chemical Information and Computer Sciences, 39(3), 525–533. 10.1021/ci980211b

106. Zaslavsky, M. E., Craig, E., Michuda, J. K., Sehgal, N., Ram-Mohan, N., Lee, J. Y., Nguyen, K. D., Hoh, R. A., Pham, T. D., Roltgen, K., Lam, B., Parsons, E. S., Macwana, S. R., DeJager, W., Drapeau, E. M., Roskin, K. M., Cunningham-Rundles, C., Moody, M. A., Haynes, B. F., . . . Boyd, S. D. (2025). Disease diagnostics using machine learning of B cell and T cell receptor sequences. Science, 387(6736), eadp2407. 10.1126/science.adp2407

107. Zhao, Y., He, B., Xu, F., Li, C., Xu, Z., Su, X., He, H., Huang, Y., Rossjohn, J., Song, J., & Yao, J. (2023). DeepAIR: A deep learning framework for effective integration of sequence and 3D structure to enable adaptive immune receptor analysis. Sci Adv, 9(32), eabo5128. 10.1126/sciadv.abo5128

108. Zimmerman, J. M., Eliezer, N., & Simha, R. (1968). The characterization of amino acid sequences in proteins by statistical methods. J Theor Biol, 21(2), 170–201. 10.1016/0022-5193(68)90069-6

