## Supplemental File 1 for "AbSolution: interactive exploration of sequence-derived features in AIRR-seq repertoires"

### AbSolution

Rodrigo García-Valiente

**2 Reproducibility of AbSolution** **10**

### 1 A guide to `AbSolution`

`AbSolution` is an interactive tool for exploring immune repertoires and their sequence-based features. `AbSolution` has been designed under the principles of accessibility, scalability, flexibility, interactivity and reproducibility to be approachable to all users while customizable for the different AIRR-Seq studies.

This vignette offers an overview of `AbSolution`'s functionality, guides you on how to initialize the app with your own data, and how to customize the app.

#### 1.1 Launching `AbSolution`

This is easy. Only two lines are required.

```
# Load required package
library(AbSolution)

# Load Shiny app
AbSolution::run_app()
```

AbSolution.R

Make sure you have space in your hard-drive! It can very easily occupy quite a lot of gigabytes.

##### 1.1.1 Quick Help

Need customized information about your **current step**? Check the **Help subtab** in the `AbSolution` tab.

Looking for an **in-depth explanation of an option**? Activate **tooltip mode** by clicking the "?" at the top right corner to see explanations for each option when you hover over them.

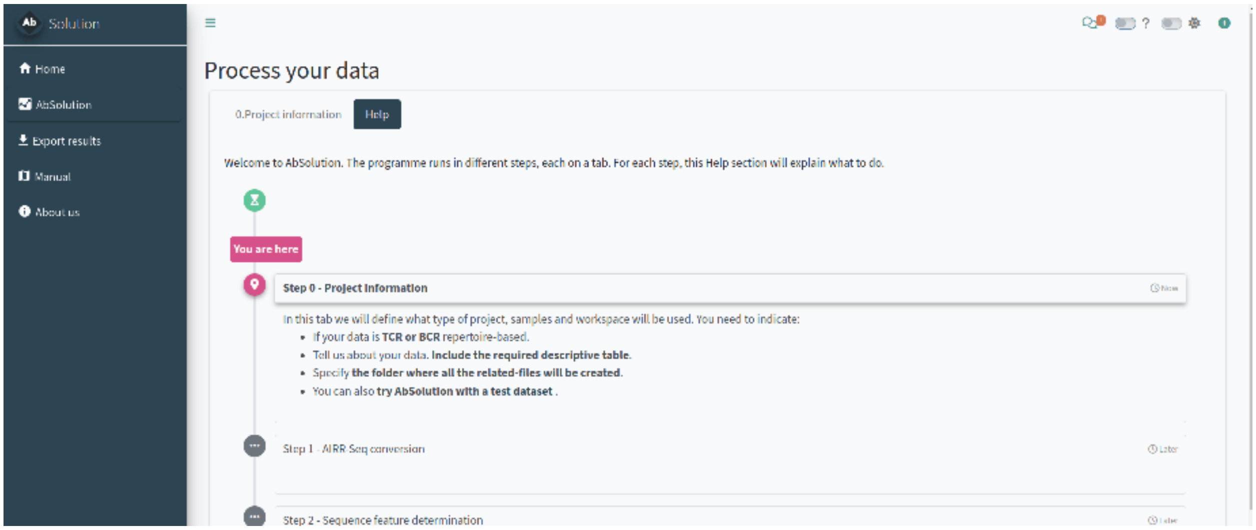

Figure 1: The Help subtab adjusts its content based on the current step the user is in.

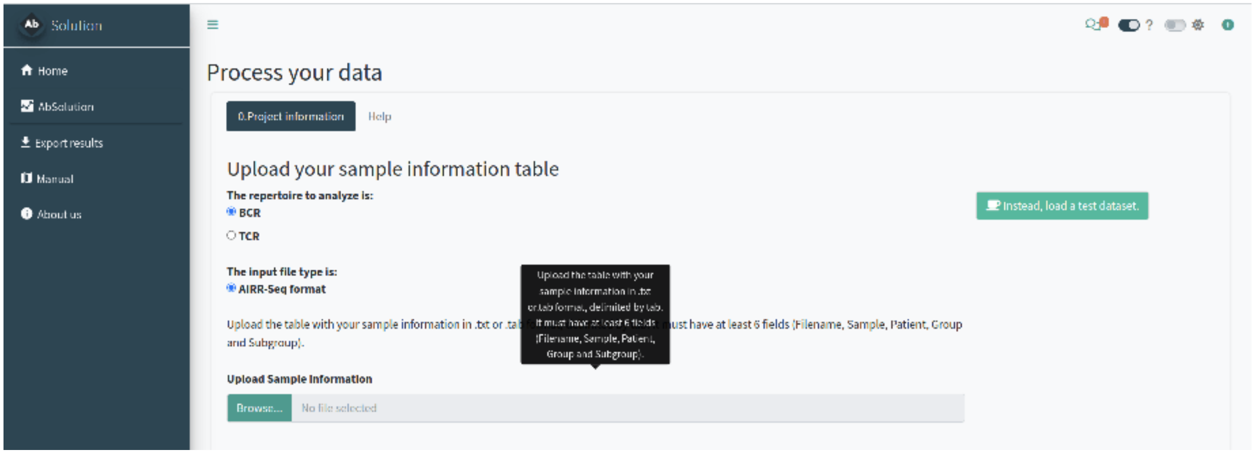

Figure 2: The tooltip mode is activated (top right corner) and a tooltip is shown over the hovered option.

##### 1.1.2 Setting up the project analysis - Step 0

In the *0.Project information* tab, we will define the type of project, samples, and workspace to be used. You need to indicate:

- If your data is **TCR or BCR** repertoire-based.
- Provide details about your data, including the **required descriptive table**.
- Specify **the folder where all related files will be created**.

##### 1.1.3 *OPTIONAL: try the test dataset*

You can **try `AbSolution` with a test dataset**. This is a small, down-sampled dataset of 10x Genomics Ig V(D)J sequences from CD19+ B cells isolated from the PBMCs of a healthy human donor. The data,

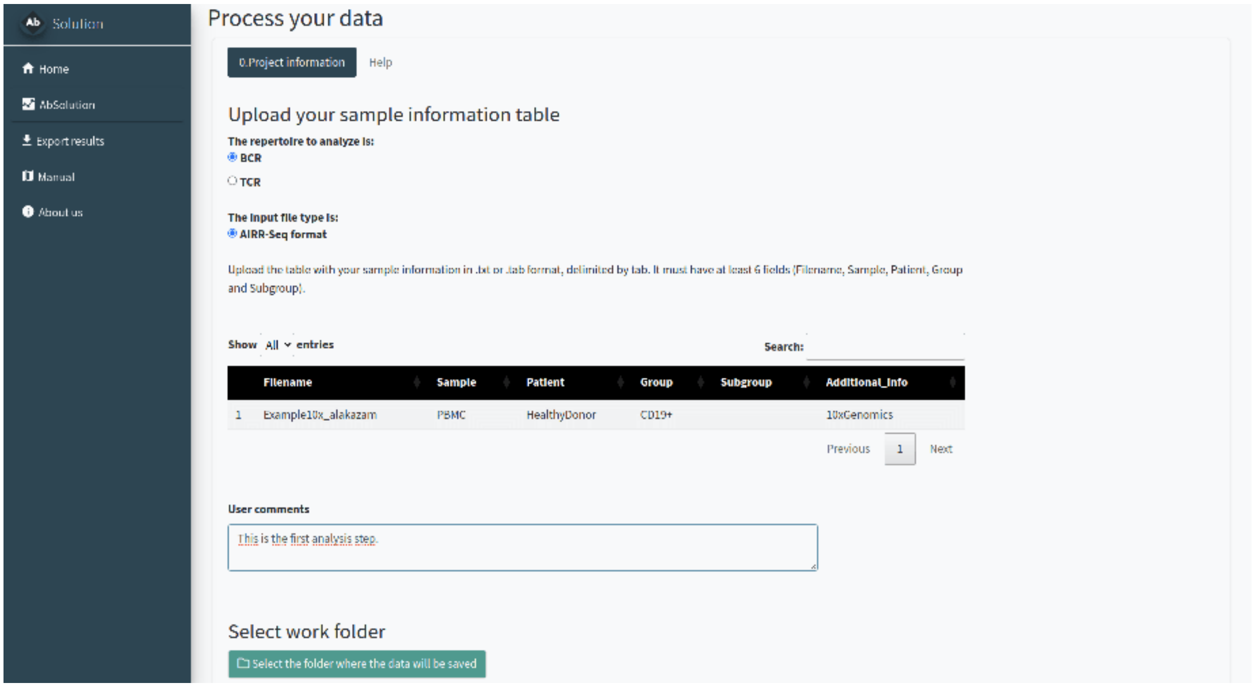

Figure 3: Tab 0.Project_information. Setting up the project analysis.

provided by 10x Genomics under a Creative Commons Attribution license, has been processed using their Cell Ranger pipeline. It comes from the Alakazam package.

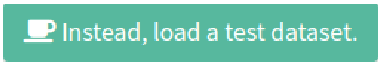

Figure 4: Button - Load the test dataset.

Just click the *Instead, load a test dataset* button and continue the workflow.

##### 1.1.4 Parsing the sequences - Step 1

In the *1.AIRR-Seq conversion* tab, we will reconstruct the repertoire germlines based on your input. `AbSolution` works with full Fab sequences or sequences with partially-sequenced FWR1/4. You need to indicate:

- The **folder where your data is located**. All files must be in the same folder. Filenames should match those indicated in the previous step. If they are found in the folder, it will say 'Yes' in Found_in_folder; otherwise, it will say 'No'.
- Additional information to better **reconstruct the germlines**. Decide whether to keep the CDR3 D-gene segment as is or use the D gene information to reconstruct it. Indicate if the FWR1 or FWR4 are partially sequenced and if there is a pre or post-Fab sequence to be removed.

Please confirm with the pre-visualization that the sequences are correctly parsed and reconstructed. A lowercase letter in the repertoire sequence indicates an insertion, while a lowercase letter in the reconstructed germline indicates a deletion in the repertoire.

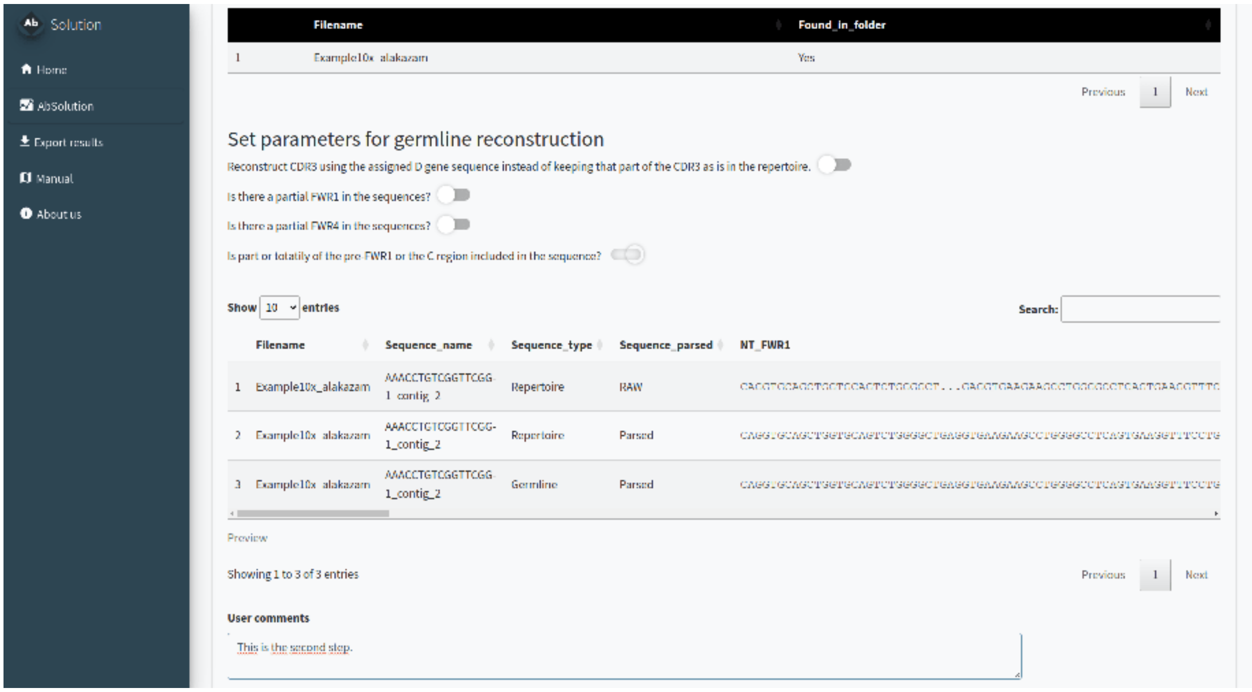

Figure 5: Tab 1.AIRR-Seq conversion. Parsing sequences and reconstructing germlines.

###### 1.1.5 Calculating sequence-based features - Step 2

In the *2.Sequence feature determination* tab, we will calculate the features (physicochemical, composition, etc.) at the nucleotide (NT) and amino acid (AA) levels for the entire sequence (repertoire and reconstructed germline) and its individual regions.

Just click the button, and wait until it is completed. It can take quite some time depending on your dataset size!

###### 1.1.6 *ALTERNATIVE: start with previously calculated files - Skip steps 1 & 2*

For pre-calculated data, in step 0 you can select to skip steps 1 and 2. In the **1&2.Select your FBM and associated files** tab, you can load the files and proceed directly to step 3.

You need to specify the **folder where your pre-calculated FBM files are located**. All files must be in the same folder, and filenames should match those indicated in the previous step. If the files are found in the folder, it will display ‘Yes’ in the Found_in_folder column; otherwise, it will display ‘No’.

You can also generate dummy datasets with the same number of mutations to use as a negative reference in the next steps.

###### 1.1.7 Exploring the dataset and filtering sequences and variables - Step 3

In the **3.Dataset exploration and variable selection** tab, you can customize your analysis by selecting specific sequences, variables, and groups, ensuring a tailored and comprehensive exploration of your dataset. Variables with NAs, infinite values, or zero variance are automatically removed.

There are different steps and options inside:

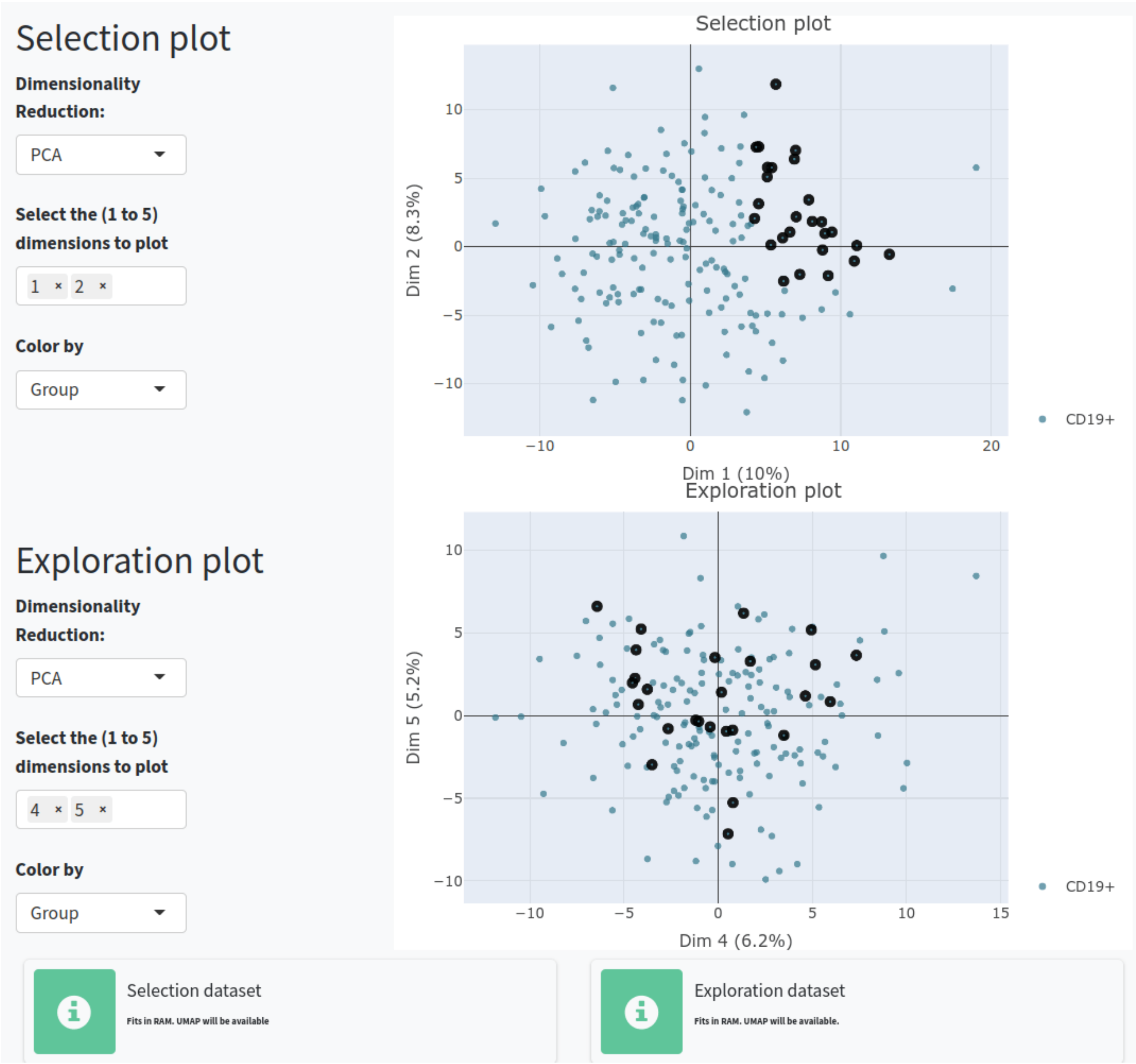

Figure 6: Tab 3.Dataset exploration. PCA visualization of BCR sequences according to the filtered variables. The variable was selected after differential analysis between a manually selected group against the rest of the sequences. Sequences in the Selection set will be shown in the **Feature Exploration** and **Clonal Exploration** tabs. The Exploration set can be used to try combinations and compare with the Selection set. Selected sequences are highlighted in both visualizations and positions and behaviours can be compared.

1. **Select and filter the sequences.** You can choose whether to include repertoire sequences, reconstructed germline sequences, or both. You can also filter sequences based on productivity, sample, chain type, and the number of replacement (R) mutations.

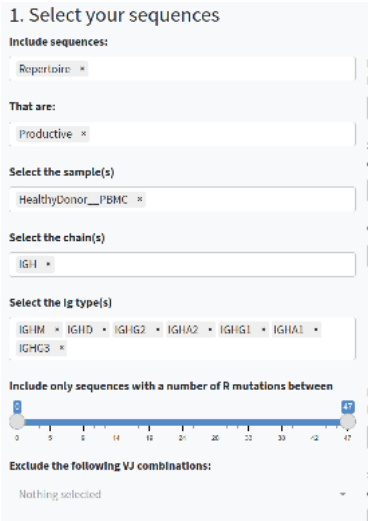

Figure 7: Options to select and filter sequences.

2. **Select and filter the variables.** You can choose which regions (e.g., CDR3, FWR1, Whole Fab), types of sequences-based features (e.g., AA-based, NT-based, composition, physicochemical, etc.) to include in the analysis, or exclude specific features. You can also work with the baseline values or with the germline-difference values.

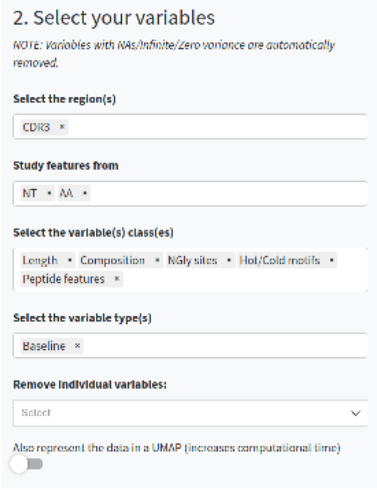

Figure 8: Options to select and filter variables.

3. **Decide if you want to compare between groups of sequences.** If so, specify which sequences and the type of comparison you want to perform. You can specify groups of sequences for comparison and normalize groups per VDJ usage or clonal usage to avoid overrepresentation. You can also apply these rules to individual samples instead of at a group-level and include only features correlated to the groups.

4. **Perform calculations** for the Selection set (which will be shown in the **Feature Exploration** and **Clonal Exploration** tabs) or the Exploration set (which you can use to try combinations and compare

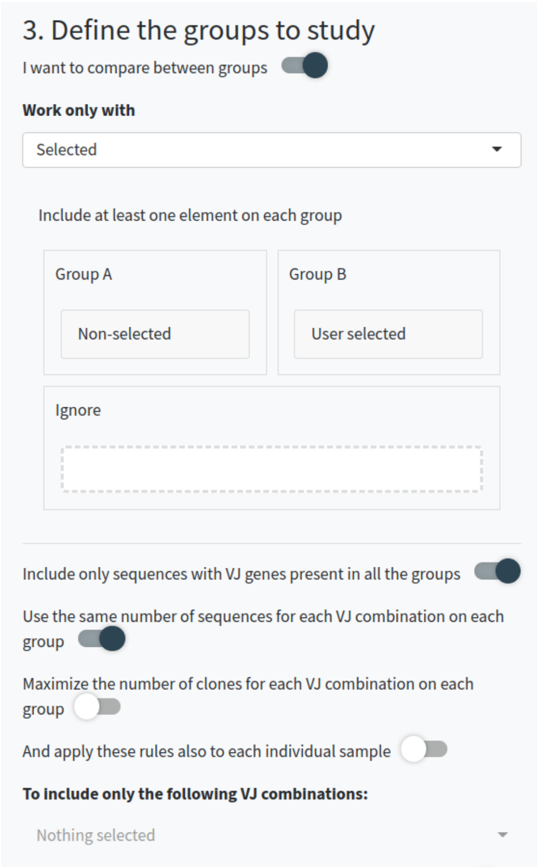

Figure 9: Options to compare between groups.

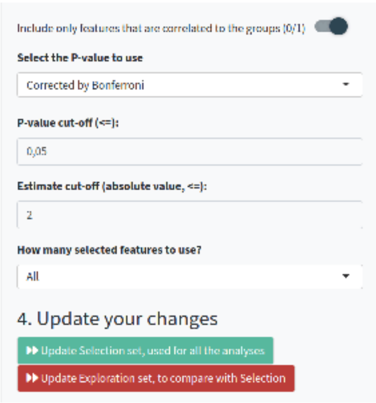

Figure 10: additional options to compare between groups and action buttons to perform calculations on the Selection or Exploration set.

with the Selection set).

##### 1.1.8 Exploring the variables - Step 4

In the *4.Feature exploration* tab, you can plot the filtered variables as a violin plot to observe their behavior according to the selected group in field 4 of the **Dataset Exploration and Variable Selection** tab. You can add the values as dots, select those of interest, and include the reconstructed germline sequences (if they are not already included) to compare how these values have changed since the germline sequence.

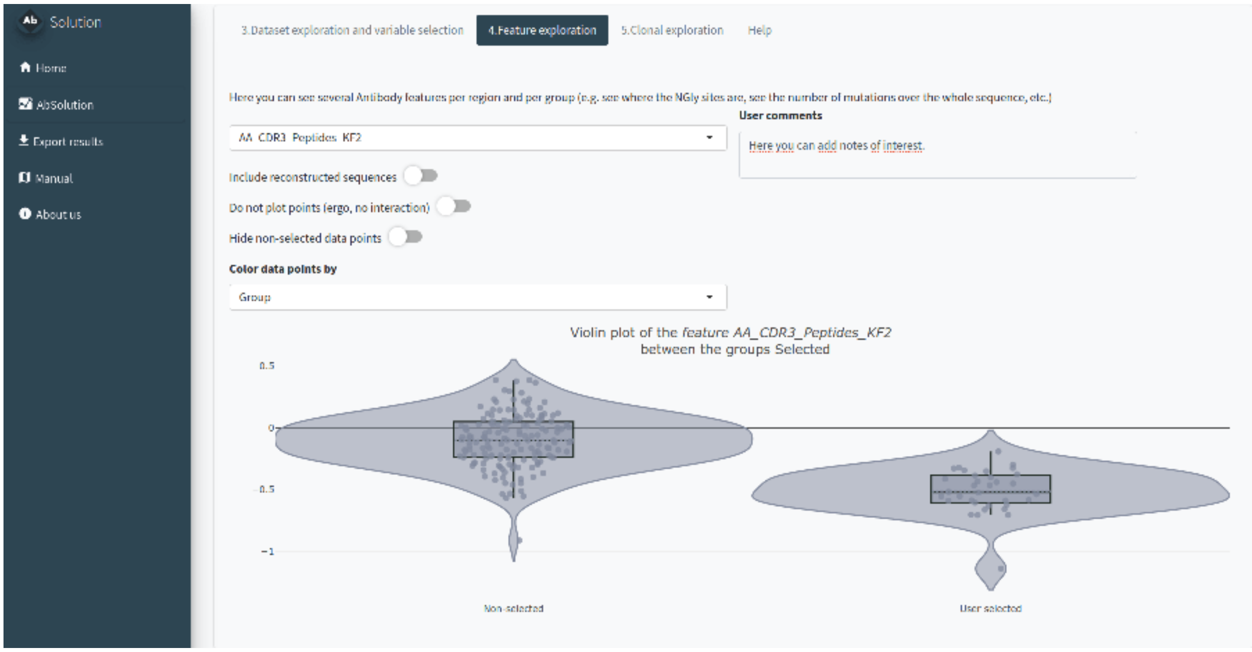

Figure 11: Tab 4.Feature exploration. Options and violin plots displaying the Kidera Factor 2 CDR3 values for two BCR groups. The variable was selected after differential analysis between a manually selected group against the rest of the sequences.

##### 1.1.9 Exploring the clones - Step 5

In the *5.Clonal exploration* tab, we will define the clonal definition to be used. You can:

- Use a pre-existing clone definition (Clone_ID) or calculate it de novo.
- Select a dominance threshold based on relative abundance.
- Identify shared clones between samples.
- Manually select clones of interest.

If you have a clonal definition assigned and the images in the clonal and feature tabs have loaded, you will be able to export results in the **Export Results** tab.

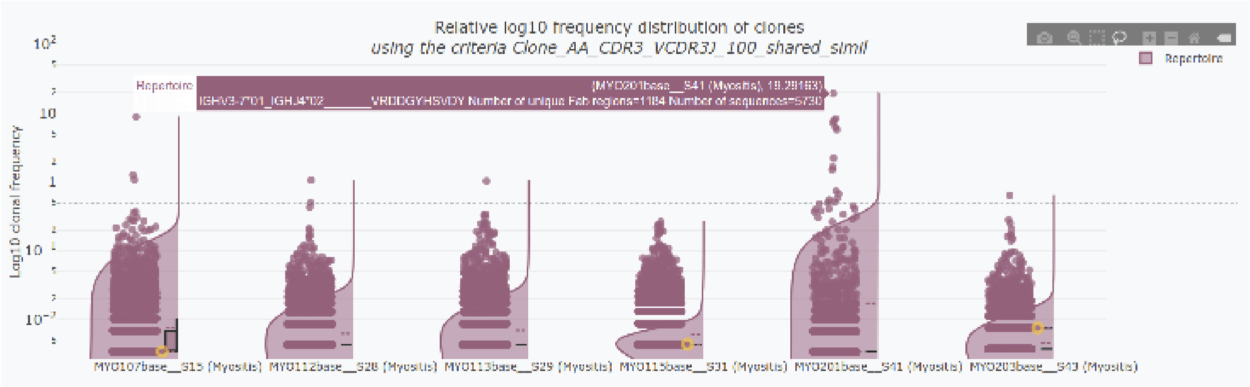

Figure 12: Tab 5.Clonal exploration. Violin plots displaying the log10 relative frequency distribution of BCR clones across six samples. Highlighted in yellow, a shared clone between samples. The horizontal dashed line represents the dominance threshold. On the top, information regarding clone composition of a hovered clone.

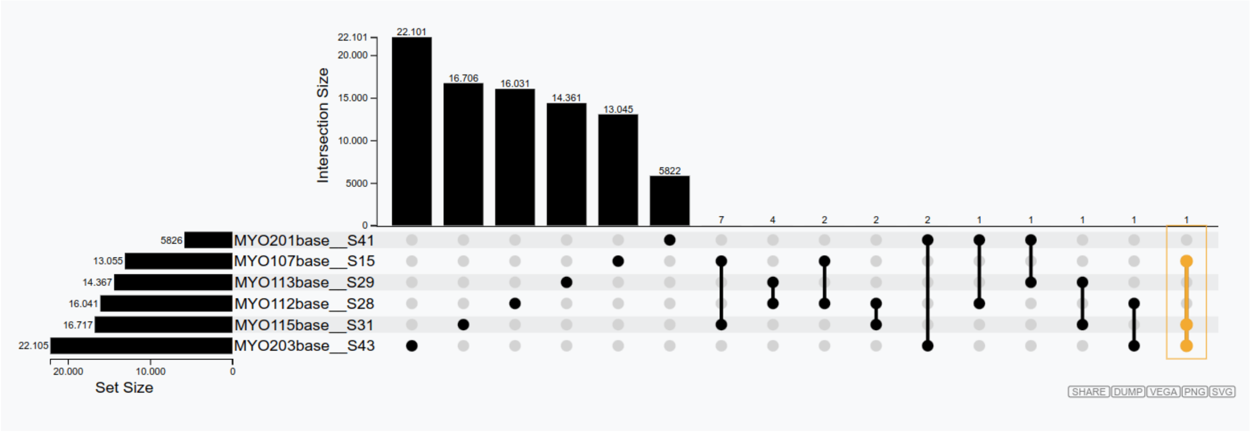

Figure 13: Tab 5.Clonal exploration. Upset plot displaying the shared BCR clones across six samples. Highlighted in yellow, a shared clone between samples. The selected clone is highlighted accordingly in the other plots.

#### 2 Reproducibility of `AbSolution`

In the *Export results* tab, after running a full analysis with the app (this means you need to have generated clonal figures in Step 5), you can download a .zip file that follows the ENCORE structure. This .zip file contains the following subfolders:

- **/0__SoftwareEnvironment/R/**: Contains the `AbSolution` package in the version used for the analysis, a Docker file, and the renv information to reproduce everything.
- **/Data/Dataset/**: Contains the dataset used for the analysis.
- **/Raw/**: Contains the sample input files.
- **/Meta/**: Contains a file with information about the samples.
- **/Processed/**: Contains the files produced during the analysis with `AbSolution` (sequence and feature information). These files are used for analysis and data exploration.
- **/Notebook/**: Contains the .Rmd file to generate the figures. This .Rmd file is generated directly from the app's code, capturing the domain logic using shinymeta. Additionally, once exported, it uses relative paths instead of absolute paths, making it easy to knit successfully. The chunks `parse_input` and `feature_calculation` are not evaluated unless the user sets `eval=TRUE`, as they are used to produce the Processed files. For time efficiency, this part is skipped when producing the .zip. There are various fields along the pipeline where users can take notes. These notes will be included in the subsequent notebook and results.
- **/Results/**: Contains the HTML file produced from the .Rmd file with the key plots from `AbSolution`. The HTML file includes all the system information and package versioning details.

It can take some time to load and some time to generate this .zip file. Be patient!
